# Global and tissue-specific aging effects on murine proteomes

**DOI:** 10.1101/2022.05.17.492125

**Authors:** Gregory R. Keele, Ji-Gang Zhang, John Szpyt, Ron Korstanje, Steven P. Gygi, Gary A. Churchill, Devin K. Schweppe

## Abstract

Maintenance of protein homeostasis degrades with age, contributing to aging-related decline and disease. Previous studies have primarily surveyed transcriptional changes with age. To define the effects of age directly at the protein level, we performed discovery-based proteomics in 10 tissues from 20 C57BL/6J mice, representing both sexes at adult and late midlife ages (8 and 18 months). Consistent with previous studies, age-related changes in protein abundance often have no corresponding transcriptional change. Aging resulted in increases in immune protein abundance across all tissues, consistent with a global pattern of immune infiltration with age. Our protein-centric data revealed tissue-specific aging changes with potential functional consequences, including altered endoplasmic reticulum and protein trafficking in the spleen. We further observed changes in the stoichiometry of protein complexes with important roles in protein homeostasis such as the CCT/TriC complex and large ribosomal subunit. These data provide a foundation for understanding how proteins contribute to systemic aging across tissues.

## INTRODUCTION

Aging results in a progressive decline in physiological function that increases risk of disease and ultimately death^1^. Gene expression studies have revealed age-related changes in transcripts that are shared across tissues, and others that are tissue-specific^2–4^, as well as transcripts that contrast healthy and diseased aging in humans^5^. However, these studies are unable to directly ascertain age-related changes in proteins. Untargeted, quantitative proteomics studies can reveal how proteins change with age, which can confirm findings from gene expression, but more importantly, they can identify molecular aging signatures that occur independently of gene expression changes.

We have previously investigated the effects of age and sex on gene expression and protein abundance in kidneys^6^ and hearts^7^ from genetically diverse mice. We found that differences in protein abundance between males and females in the kidney were largely mediated through their transcripts. In contrast, differences in protein abundance across ages were largely independent of their transcripts. A similar dynamic between sex and age differences was observed in heart, although fewer sex differences were present. From these studies, we concluded that many age-related changes in protein abundance are not due to corresponding changes in gene expression and that transcriptomics provides an incomplete picture of aging in kidney and heart. Multiple mechanisms could result in decoupling proteins from their transcripts with age, including reduced proteasome activity^8^ and reduced ribosome occupancy^9^ with age.

A key question is whether this discordance between protein abundance and gene expression occurs for most tissues. Furthermore, we are interested in which age-related changes to proteins are shared across tissues and which are unique. Comparing the kidney and heart revealed common signatures of increased immune cell infiltration in both tissues for proteins and transcripts. We also observed tissue-specific changes, particularly among proteins. Changes in the kidney proteome correspond to functions specific to the substructures and cell subtypes of the nephron, including the podocytes and proximal tubule cells. In the heart, we observed age-related changes in fatty acid metabolism and autophagy. These tissue-specific changes relate to the unique biological functions and stressors of these tissues during aging.

To address these questions, we performed a survey of protein abundance across ten tissues (kidney, liver, fat, spleen, lung, heart, skeletal muscle, striatum, cerebellum, and hippocampus) collected from female and male C57BL/6J mice at 8 and 18 months of age. We performed multiplexed, quantitative mass spectrometry on bulk tissue samples and analyzed differences in protein abundance between age and sex, as well as sex-specific changes with age. We compared age and sex differences for proteins in our study to transcript changes in corresponding tissues that were reported by Schaum *et al* 2020^2^. We used enrichment analyses to assess how aging affects biological processes, as reflected by coordinated changes in proteins across and within tissues. Finally, we characterized aging-related changes to protein complexes, in terms of overall and relative abundances of member proteins. Our findings confirm that the discordance between age-related changes in proteins and gene expression occurs across multiple tissues. Our data survey a broad range of age-related changes in proteins that occur globally across tissues and others that are tissue specific.

## RESULTS

We quantitatively profiled protein abundance across 10 tissues (Table S1) representing a range of organ systems (kidney, liver, fat, spleen, lung, heart, skeletal muscle, striatum, cerebellum, and hippocampus) from 20 C57BL/6J (B6) mice using tandem mass tags (TMT) and real-time search (RTS) mass spectrometry^10^. Animals represented an equal balance across males, females, and two age groups (8 and 18 months), with five animals per age-by-sex group. Outlying samples were identified using principal component analysis (PCA), resulting in the removal of one sample from liver, fat, spleen, lung, and skeletal muscle, and two samples from striatum (Methods). Outlying samples across tissues were not from the same mouse. Cumulatively, we detected 11,853 proteins across the 10 tissues. We filtered the data to a high confidence set of proteins (observed in both batches per tissue), resulting in 10,250 proteins used in further quantitative analysis (Figure 1, Figure S1A). Spleen had the highest number of analyzed proteins (6,556) and skeletal muscle had the fewest (2,353) (Upset plot^11^ in Figure S1B). Many proteins were detected across multiple tissues. We observed 676 different protein-to-tissue overlap sets, with detection in all tissues being the most common (1,229) followed by proteins detected in only spleen (1,003) and then only the three brain tissues (425) (Figure S1B).

**Figure 1.**
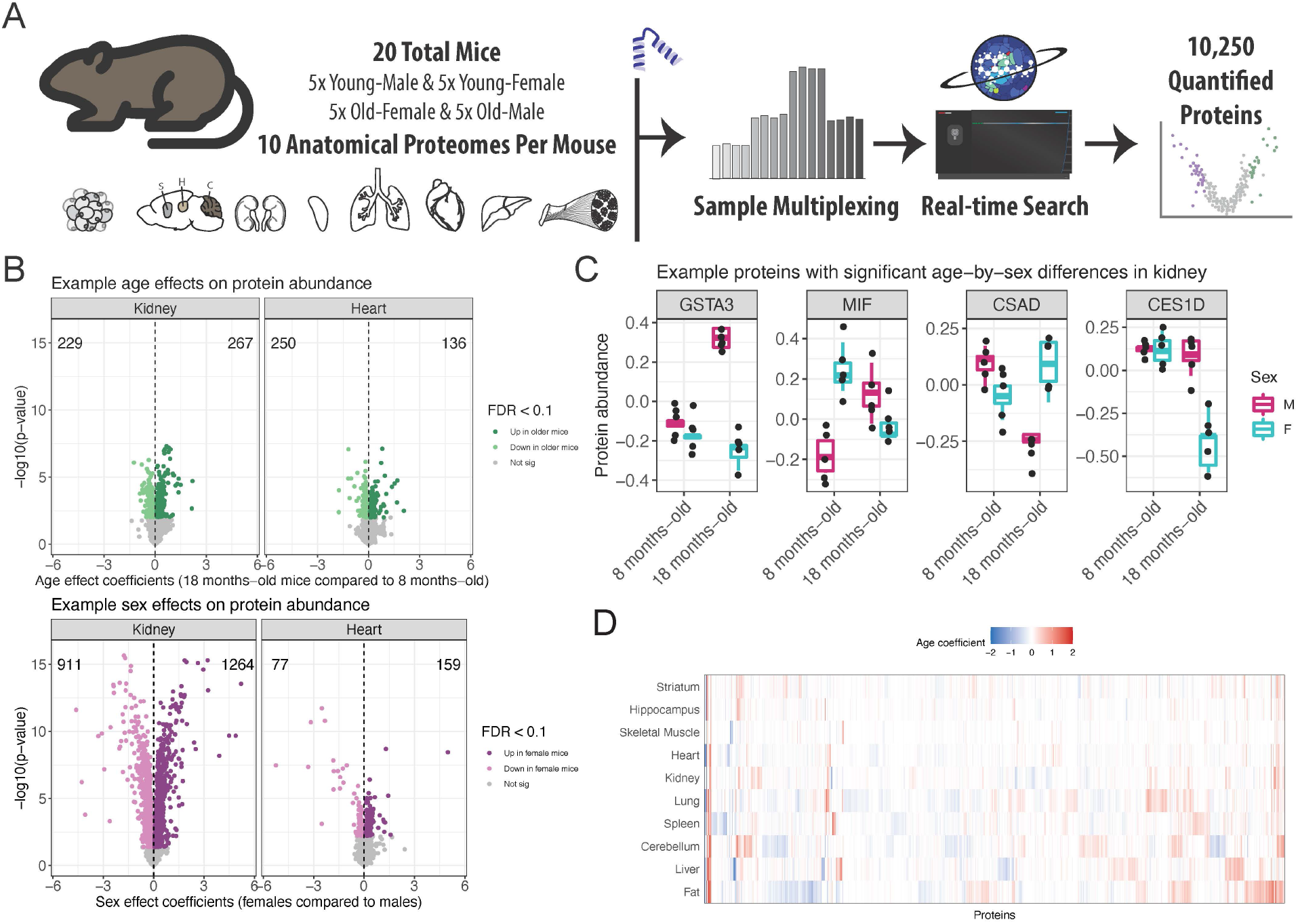
Quantitative proteomics study on the effects of age and sex on protein abundance across 10 tissues from B6 mice. (A) Using sample multiplexing, 10 anatomical proteomes (adipose tissue, striatum, hippocampus, cerebellum, kidney, spleen, lung, heart, liver, skeletal muscle) were profiled across age and sex matched mice (n=20). (B) The top 25 intersections of proteins and the tissues in which they were detected and analyzed. (C) Age (top) and sex (bottom) differences for protein abundance from kidney (left) and heart (right) tissues, depicted as volcano plots. Differences in protein abundance are summarized as regression coefficients (x-axis) and corresponding −log_10_(*p*-value) (y-axis). Points are colored based on statistical significance (FDR < 0.1) and direction of effect. Counts of proteins with significantly higher abundance in 18 month-old mice and 8 month-old mice are included. Dashed vertical lines at 0 included for reference. (D) Examples of proteins in kidney tissue with significant age-by-sex differences (FDR < 0.1). (E) Age differences detected across the 10 tissues (FDR < 0.1), represented as a heatmap. Differences are summarized as regression coefficients.

To assess how different technical features of the experiment contributed to variation in the abundance of individual proteins, we jointly modeled data across tissues by fitting random effects models (Methods) for each of the 1,229 proteins measured in all 10 tissues (Table S2). As expected, the protein abundance varies greatly with tissue/batch, which are confounded due to each tissue being measured in two separate runs, *i.e*., batches, of the mass spectrometer (Figure S1C). We note that our goal here is not to detect protein abundance differences between tissues, which would be invalid due to confounding, but rather to detect patterns in how abundance changes with age across tissues. We identified 13 proteins that had abundance patterns that were highly specific to an individual mouse across the 10 tissues, including IGHG2C and four other immunoglobulins (Figure S1C-F). Using gene set enrichment analysis^12^ (Methods), the 13 proteins (compared to the overall set of 1,229 proteins observed in all tissues) were enriched for gene ontology (GO) terms related to adaptive immune response, which suggests that the mouse may have been experiencing an active infection.

### Effects of age and sex on the abundance of individual proteins

Protein abundance can vary with age^6,7,13^ and between sexes^14–16^. For each tissue, we characterized age and sex effects (Table S3) and declared differences to be significant based on false discovery rate (FDR) < 0.1 (Figures 2 and S2, respectively; Methods). The number of proteins with age effects varied greatly across the tissues, ranging from lung with the most (866) to striatum with none meeting statistical significance (partially due to loss of power from the removal of two animals). The number of proteins with sex effects varied across the tissues, from kidney with 2,175 to the three brain tissues with 10 or fewer. We also identified proteins for which sex differences in abundance depended on age by testing for age-by-sex interaction effects (Methods), detecting 21 in kidney, 6 in liver, 5 in fat, and 11 in skeletal muscle (FDR < 0.1) (Figure S3). The skeletal muscle proteins, for which males had distinctly higher abundance than females in older mice, were associated with the endoplasmic reticulum lumen. Comparison of sex and age effects across the tissues revealed generally more differences between sexes than between ages, most notably in kidney, liver, fat, and spleen. However, proteins with significant age effects outnumbered those with sex effects in lung, heart, and cerebellum (Figure S4). The three brain tissues were distinctly buffered from differences based on age and sex. In total, 2,356 proteins had a significant age effect in at least one tissue, 5,125 had a sex effect, and 43 had an age-by-sex interaction effect (FDR < 0.1).

**Figure 2.**
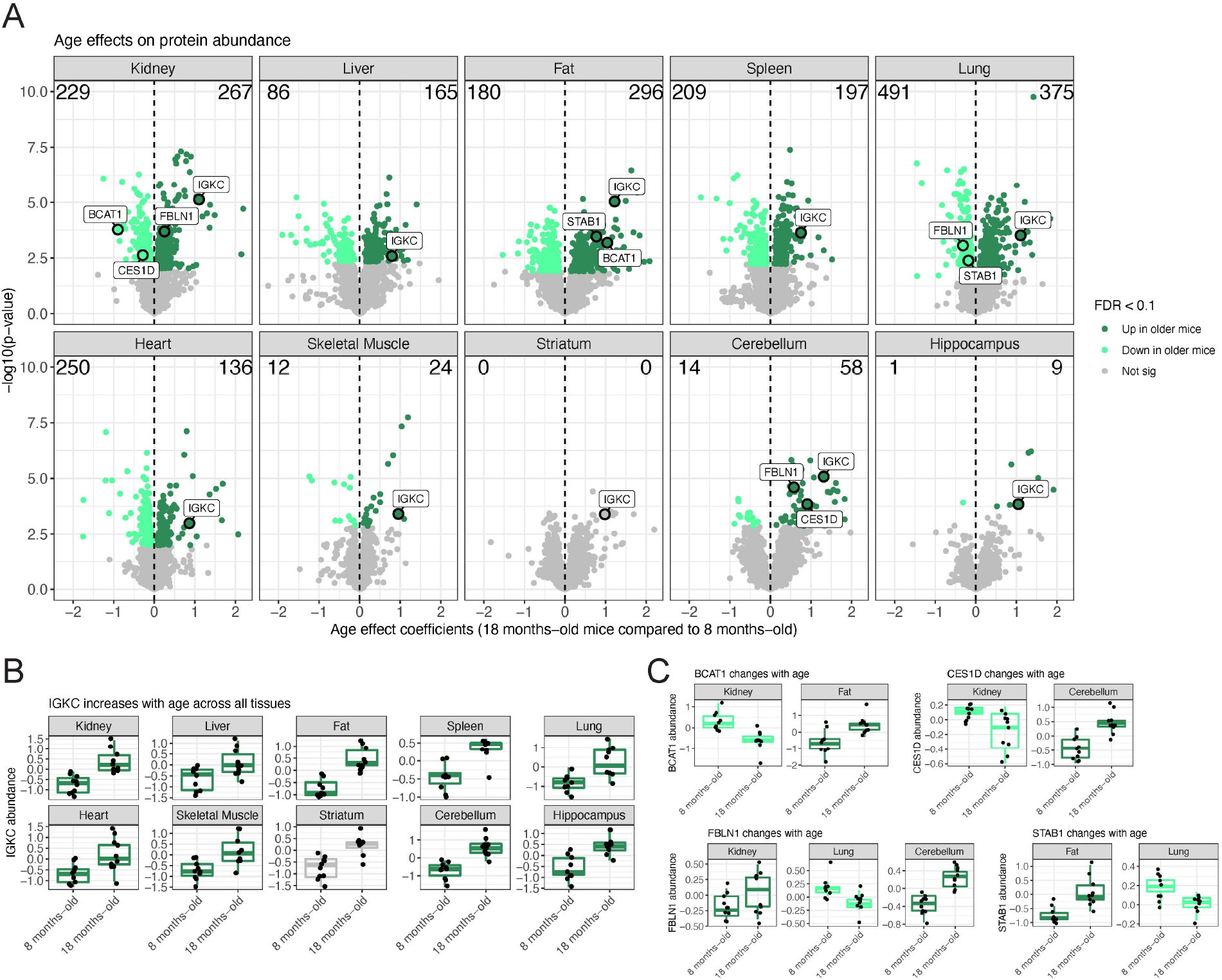
Proteins with age-related differences in abundance, across 10 tissues. (A) Proteins with age differences in abundance, represented as volcano plots. Differences in protein abundance are summarized as regression coefficients (x-axis) and corresponding −log_10_(*p*-value) (y-axis). Points are colored based on statistical significance (FDR < 0.1) and direction of effect. Counts of proteins with significantly higher abundance in older mice and younger mice are included. Dashed vertical lines at 0 included for reference. Proteins with sex differences in abundance are shown in Figure S2. (B) The immunoglobulin IGKC has consistent increased abundance in older mice across all 10 tissues. (C) Proteins with significant age differences that vary between tissues: BCAT1, CES1D, FBLN1, and STAB1.

We jointly modeled individual proteins across multiple tissues (Methods) to test whether the age or sex effects on proteins were consistent across tissues or unique to specific tissues (Table S4), declaring significance based on FDR < 0.1. Among the 7,745 proteins detected in more than one tissue, 643 had consistent age differences across the tissues in which they were quantified. For example, IKGC is an immunoglobulin that has increased abundance in older mice for all tissues (cross-tissue age *p* = 1.1e-6; Figure 2B). We detected 1,028 proteins with age effects that differed between tissues, representing proteins that had age differences in only some of the tissue or even age effects with differing directions, such as BCAT1 (age-by-tissue *p* = 1.8e-10), CES1D (age-by-tissue *p* = 1.1e-10), FBLN1 (age-by-tissue *p* = 2.1e-8), and STAB1 (age-by-tissue *p* = 2.6e-8) (Figure 2C). These age-related differences reflect tissue-specific features related to aging decline. For example in kidney, there was reduced abundance of BCAT1, which promotes mitochondrial biogenesis and ATP production and has been shown to promote breast cancer formation when knocked down^17^, and REN1, which has been shown to play a role in modulating vascular tone and tubular function in the kidney^18^. For sex effects, we detected 1,006 proteins with consistent differences between sexes across tissues and 2,565 proteins with tissue-specificity.

### Most age-related changes in protein show no corresponding transcriptional change

To compare age effects on proteins to corresponding effects on transcripts, we obtained data from Schaum *et al* 2020^2^, in which bulk RNA sequencing was performed across 17 tissues of mice from 10 age groups, ranging from 1 to 27 months. Each age group consisted of 4 to 6 C57BL/6JN mice. Overlapping tissues between studies include kidney, liver, and heart. We selected the transcriptomics data from the 9 and 18 months age groups, which are closest to 8 and 18 months in the proteomics data, and characterized the age and sex effects on transcripts (Methods). We contrasted the age effects between proteins and their transcripts, and for comparison, we also compared the sex effects on proteins and their transcripts.

The age effects on proteins and transcripts are generally not consistent. In contrast, sex differences are highly concordant between proteins and transcripts, most notably in kidney and liver (Figures 3A-C). The consistency of strong sex effects correlated between protein and transcript supports the validity of comparing data across distinct but related mouse strains^19^ with mice that were raised at different sites as part of different experiments. We observed a similar dynamic between age and sex differences in the kidneys of genetically diverse mice^6^. Genes with consistent age effects on transcripts and proteins in kidney include *Keg1* and *Vcam1* (Figures 3D-E). VCAM1 is an immunoglobulin that facilitates interactions between vascular and immune cells and its increase with age has been associated with age-related disease in humans^20–22^. We observed increases in VCAM1 abundance with age across multiple tissues (kidney, liver, fat, and cerebellum), which resulted in a significant cross-tissue age effect (cross-tissue age *p* = 1.59e-7). We note that the expression data suggest greater variation within ages and sexes than the protein data, which has more mice per age-by-sex group.

**Figure 3.**
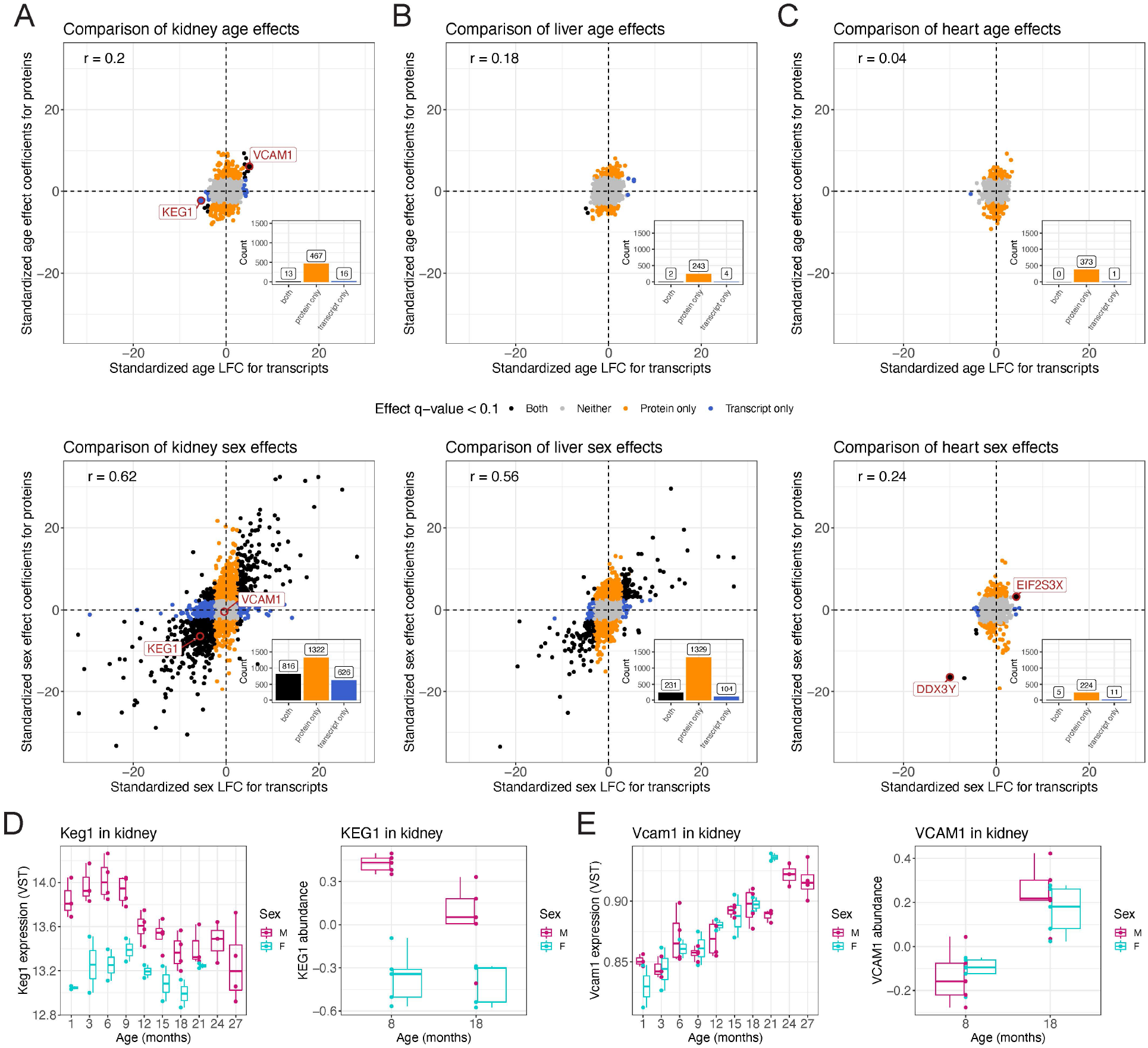
Age-related differences are less shared between protein and transcripts than sex-related differences. Comparisons of age- (top) and sex-related (bottom) differences in proteins to transcripts in (A) kidney, (B) liver, and (C) heart. Points are colored based on statistical significance (FDR < 0.1) in proteins and transcripts. Dashed vertical and horizontal lines at 0 included for reference. Counts of genes with significant differences included as bar plots in the bottom right quadrant. (D) *Keg1* expression had significant age and sex differences (left), whereas its protein had a matching sex effect (right). The age effect did not meet significance at FDR < 0.1, but the direction is consistent with transcripts. Transcript data represent 10 age groups compared to 2 age groups for proteins. (E) *Vcam1* significantly increase with age in terms of both transcripts (left) and proteins (right). Transcript data represent 10 age groups compared to 2 age groups for proteins.

Heart had the least consistency between the age effects on proteins and on their transcripts. It also had far fewer significant sex effects than kidney or liver, consistent with our previous work^7^, though they are still more consistent between proteins and transcripts than age effects. Notable genes with consistent sex effects on proteins and transcripts are *Ddx37* and *Eif2s3x* which are encoded on the Y and X chromosomes, respectively. These findings validate the quality of the protein data by demonstrating the consistency of sex effects with transcript data. They also highlight the importance of assessing aging-related changes among proteins, many of which are not observed at the transcript level.

### Immune-associated proteins change with age across all tissues

We clustered the 2,356 proteins that had significant age effects in at least one tissue after setting effects to zero in tissues in which proteins were not observed. Looking broadly across the clustered age effects reveals consistent immune-related differences between age groups across all 10 tissues (Figure 4A). Components of the innate immune system, most notably members of the complement cascade such as C8A and C8B, were less abundant in all tissues of older mice. Immunoglobins and other proteins related to humoral immunity were distinctly more abundant in older mice. Proteins involved in proteolysis, including immunoproteasomes like PSMB8, were also more abundant in older mice to varying degrees across the 10 tissues.

**Figure 4.**
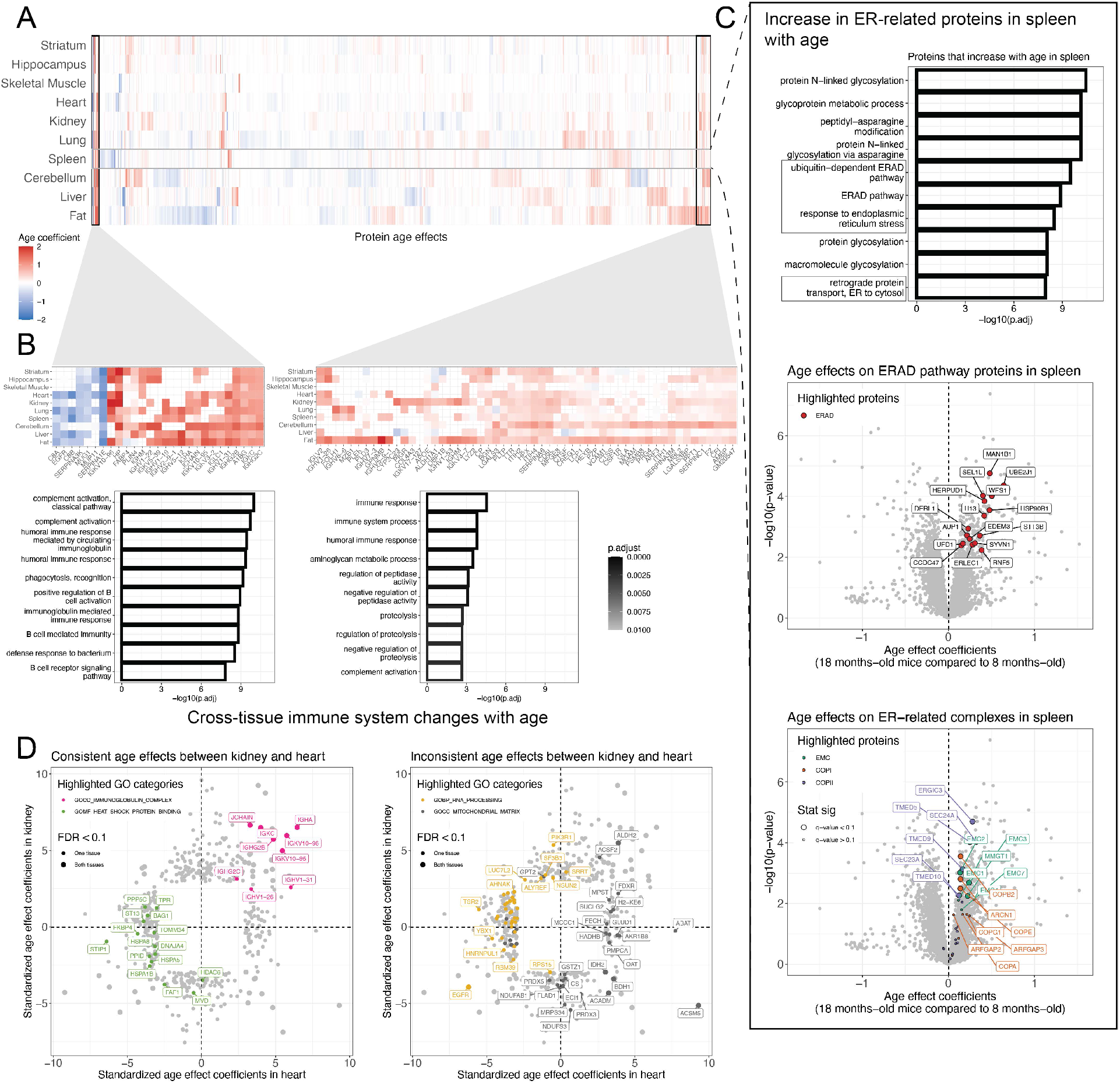
Cross-tissue and tissue-unique patterns of aging. (A) Age-related differences detected across the 10 tissues (FDR < 0.1), represented as a heatmap. Differences are summarized as regression coefficients. Hierarchical clustering of the proteins (columns) reveals sets of proteins with age difference patterns across tissues and unique to specific tissues. (B) Proteins with age differences that are shared across tissues are enriched for immune-related GO categories. Additional tissue-unique patterns are highlighted in Figure S5. (C) The proteins with age differences in a specific tissue can be enriched in GO categories, with spleen highlighted here for proteins with higher abundance in older mice. Abundance differences with age for proteins analyzed in spleen, represented as volcano plots. Differences in protein abundance are summarized as regression coefficients (x-axis) and corresponding −log_10_(*p*-value) (y-axis). The ERAD pathway, EMC, COPI, and COPII proteins are highlighted. Highlighted proteins with significant differences (FDR < 0.1) have larger point size. Proteins with age *p* < 0.05 were labeled. (D) Comparison of age differences between kidney and heart with highlighted GO categories that are consistent (left) and inconsistent (right) between the tissues. Proteins with a significant age difference (FDR < 0.1) in kidney or heart are shown. Proteins with significant differences (FDR < 0.1) in both tissues have a larger point size.

The consistency of age effects across tissues for immune-related proteins is striking, as highlighted by the adaptive immune response GO set (GO:0002250) (Figure 5A). Even tissues with few statistically significant age effects, most notably striatum with none, show differences in the same direction, such as increased abundance of immunoglobins in older mice. We note that the abundance of the immunoproteasomes (PSMB8, PSMB9, and PSMB10) is higher in older mice for the tissues with the most pronounced age-related increases in immunoglobins, fat and cerebellum (Figures 5B-C). The immunoproteasomes are inducible components that replace the constitutive components (PSMB5, PSMB6, and PSMB7) in the 20S catalytic core of the proteasome^23^. The immunoproteasome is more efficient at degrading ubiquitin-labeled proteins as antigens for presentation on MHC class I molecules, a key process in distinguishing between self and non-self in adaptive immunity^24^. However, it is not a perfect marker of immune cells, as it is also expressed by non-immune cells during inflammation^25^.

**Figure 5.**
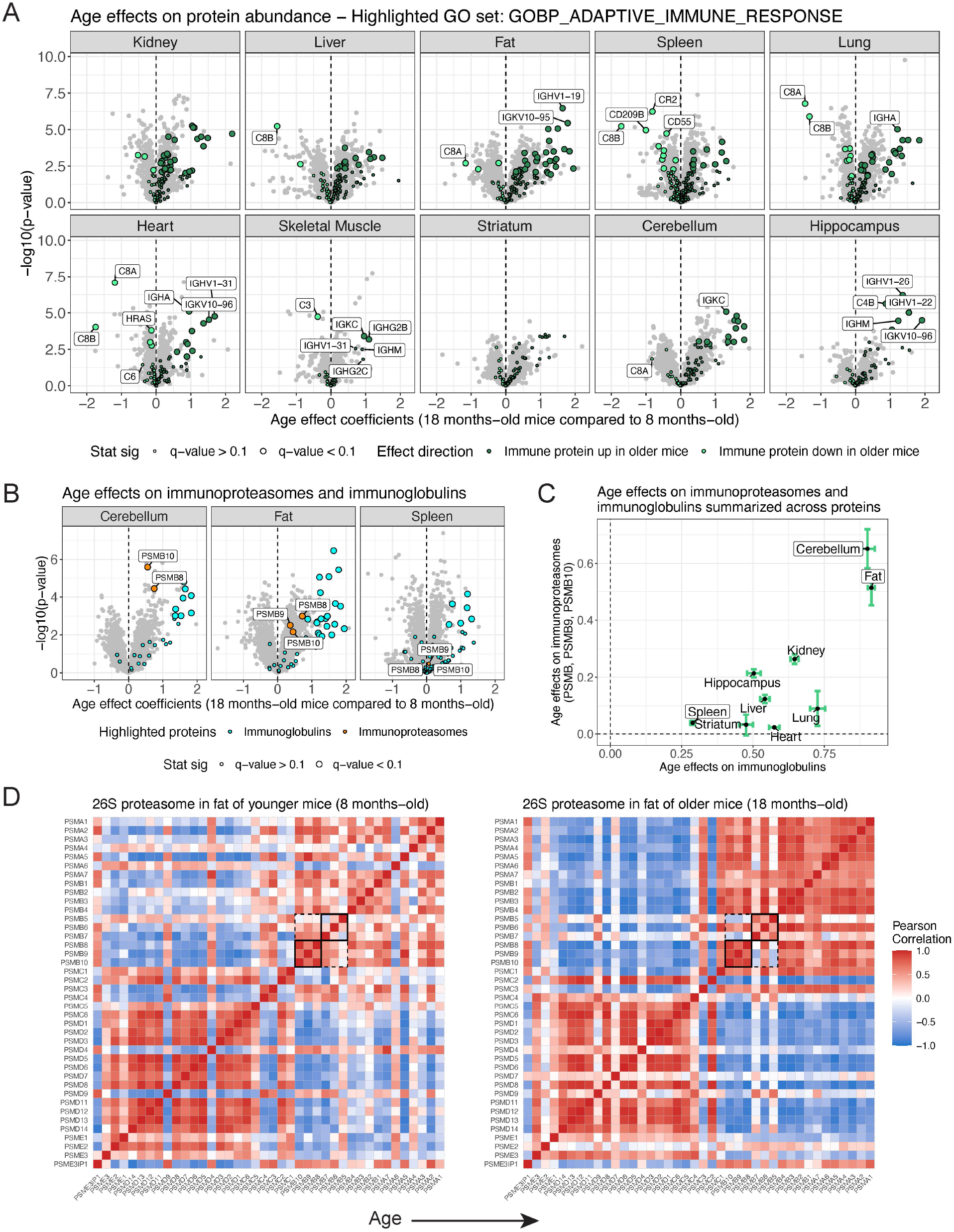
Increased immunoglobulin abundance is a signature of aging detected in all 10 tissues. (A) Proteins with age differences in abundance, represented as volcano plots. Differences in protein abundance are summarized as regression coefficients (x-axis) and corresponding −log_10_(*p*-value) (y-axis). Points are colored based on being a member of the adaptive immune response GO category and direction of effect. Highlighted proteins with significant differences (FDR < 0.1) have larger point size. Dashed vertical lines at 0 included for reference. (B) Volcano plots for cerebellum, fat, and spleen, with immunoglobins and immunoproteasomes (PSMB8, PSMB9, and PSMB10) highlighted. (C) Age differences summarized across the immunoproteasomes (y-axis) and immunoglobins (x-axis) for all 10 tissues. Points represent mean differences and bars represent standard errors. Horizontal and vertical dashed lines at 0 included for reference. (D) Pearson correlations from the proteasome in younger (top) and older (bottom) mouse fat. Rows and columns are ordered to reflect key sub-complexes of the proteasome, which are labeled. The immunoproteasomes and their matching constitutive analogs are highlighted with black square.

The co-regulation of the overall proteosome complex, encompassing the 20S catalytic core, its 19S regulator, and the 11S regulator, is multifaceted. We have previously shown that abundance of individual components and sub-complexes of the proteasome are influenced by genetic variation^15^ and age^7^ in genetically diverse mouse populations. In this B6 population where genetic variation has been fixed (excluding spontaneous mutations specific to individuals), the sub-complex structure of the proteasome, most notably the 20S catalytic core and 19S regulator, is reflected in the correlations between complex members. The individual sub-complexes become more tightly correlated within themselves (and anti-correlated with each other) in the older mice in fat tissue (Figure 5D). The correlation among the immunoproteasome components (and anti-correlation with their constitutive analogs) becomes more pronounced in the older mice, likely due to increased levels of immunoproteasome from immune cells. Changing immune cell-related tissue composition with age is further supported by the reduction in complement activation proteins with age.

### Tissue-specific signatures of aging

Age effects reveal unique patterns specific to tissues (Tables S5-6). Spleen exhibits unique increases in abundance for proteins related to the endoplasmic reticulum (ER), such as the ER-associated degradation (ERAD) pathway, ER membrane complex (EMC), and the COPI and COPII complexes (Figure 4C), suggesting potential changes to protein quality control^26^ in the spleen. The EMC enables the biogenesis of multipass transmembrane proteins and has been associated with pleiotropic phenotypes across organisms^27,28^. COPI and COPII are involved in trafficking proteins between the ER and Golgi^29^. In addition to the immune signatures observed across all 10 tissues, we saw tissue-specific immune patterns, such as decreased abundance for proteins involved in leukocyte and lymphocyte activation in the spleens of older mice (Figure S5B) and increased abundance for proteins involved in a broad range of immune system-related GO categories in the fat of older mice (Figure S5C). We observed decreased abundance in older mice for proteins related to the mitochondrial chain complex I and the oxidation-reduction process (Figure S5A) and increased abundance in older mice for proteins related to multiple metabolic processes (Figure S5D). Comparison of gene set enrichment results between tissues can highlight shared or distinct aging processes. For example, we compared kidney and heart, which revealed consistent GO categories like immunoglobulins (increasing with age) and heat shock proteins (decreasing with age) and inconsistent GO categories like the mitochondrial matrix and RNA processing (Figure 4D).

### Age and sex influence the abundance and stoichiometric balance of protein complexes

Individual members of protein complexes are often co-regulated to maintain stoichiometric balance of components^30–34^. Biological factors can affect this balance, including genetics, sex, and age^7,14–16^. More than one factor can influence a complex (or its sub-complexes or individual components), as demonstrated by the effects of both genetic variation and age on the proteasome^7,15^. To measure a protein complex’s co-abundance, referred to here as cohesiveness, for each of 228 annotated protein complexes^35,36^ across the 10 tissues, we the median correlation across all pairs of complex members (Table S7; Methods). We note that the cohesiveness of a complex could reflect its stoichiometric balance as well as cell type heterogeneity. We observed some conservation of protein complex cohesiveness between tissues, most notably among the three brain tissues (*r* > 0.68, *p* < 2.2e-16; Figure S6).

We tested whether age or sex had consistent effects on abundance across the members of a protein complex (Table S7; Methods), as would be expected if the entire protein complex is changing with age or sex. Examples include the previously mentioned COPI and COPII complexes in the spleen (Figure 4C). The number of protein complexes with a multi-protein age effect varied across the tissues, ranging from lung with 29 to skeletal muscle and hippocampus with none at FDR < 0.1 (Figure S7A). More complexes had consistent sex effects across proteins, ranging from spleen with 76 to striatum and cerebellum with none at FDR < 0.1 (Figure S7B).

Changes due to age or sex on protein complexes may be more subtle than a change in mean abundance across complex members. Cohesiveness of a protein complex can vary with age or sex. We first calculated the correlations among protein complex members for each age group and then looked for changes in overall correlation patterns with a paired *t*-test (Table S7; Methods). The same approach was used for sex. We used a stricter threshold of significance (FDR < 0.01) in order to focus on only the most significant effects. The association of age with cohesiveness of complexes varied across the tissues, ranging from 35 complexes in spleen to 3 in skeletal muscle at FDR < 0.01 (Figure S8A). For sex, spleen had the largest number of complexes with changes in cohesiveness (26), and fat, hippocampus, and striatum had the fewest with 6 each, all at FDR < 0.01 (Figure S8B). Ribosomes have been shown to lose stoichiometric balance with age in the brains of killifish^8^, and we see similar signals across many of our tissues (Figure S8A), indicating that this previous finding generalizes across tissues and species.

For each complex, we counted the number of tissues for which a significant age and sex effect on abundance or cohesiveness were detected (Figure S9). The co-distribution of age and sex effects differed between effects on abundance and effects on cohesiveness. Effects on abundance were more likely to be detected in smaller subsets of tissues, whereas effects on cohesiveness represent a distinct minority of complexes with both age and sex effects on cohesiveness across many tissues, such as the chaperonin-containing T-complex (CCT-complex) and cytoplasmic ribosomal large subunit (CRLS).

### CCT-complex is more cohesive in older B6 mouse cerebellum

The CCT-complex, also known as the tailless complex polypeptide 1 ring complex (TRiC), is required for folding proteins such as acting and tubulin. The CCT-complex was significantly more cohesive in older mouse cerebellum than in younger (*p* = 2.0e-8; Figure 6). This signal is due to CCT6A, CCT2, CCT5, and to a lesser extent CCT3, being anti-correlated with other complex members in younger mice but more correlated in older mice (Figure 6B). The significance of the change of individual correlations were determined using permutations (Methods). The pattern of correlations reflects the physical structure of the CCT-complex, which is composed of two octameric rings made from eight proteins (TCP1, CCT2, CCT3, CCT4, CCT5, CCT6A, CCT7, and CCT8)^37,38^ (Figure 6E). Notably, the CCT6A and CCT2 components from the upper and lower rings are in physical contact with their matching protein. CCT5 and CCT3 are immediately adjacent to CC2 and CCT6A, respectively, in both rings. We have previously shown that genetic variation at *Cct6a* regulates other members of the CCT-complex in genetically diverse mice^14,15^. There is no complex-wide age effect on abundance (*p* = 0.87) and none of the proteins differ significantly in mean abundance between the two age groups (Figure 6B). In older mouse cerebellum, members of the CCT-complex are correlated with more non-CCT-complex members (1,364 genes with *r* > 0.75 in older mice and *r* < 0.25 in younger mice), which enrich for many GO categories related to its function in folding cytoskeleton proteins^39^, such as the microtubule category (GO:0005874) (Figures 6F & S10).

**Figure 6.**
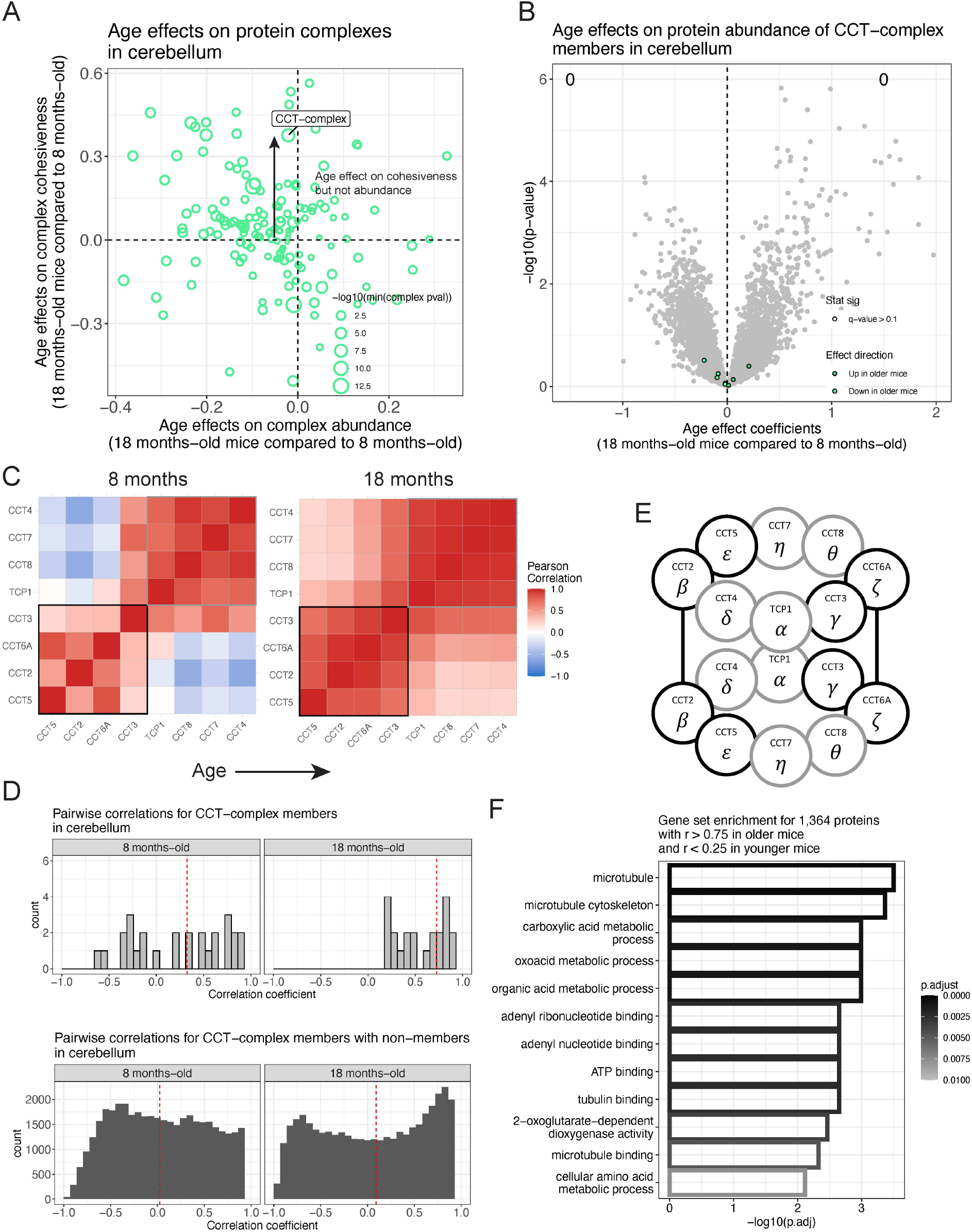
CCT-complex is more stoichiometrically balanced in older mouse cerebellum. (A) Comparison of age-related differences in complex-wide correlation (y-axis) to complex-wide abundance (x-axis) in cerebellum. Point size corresponds to the minimum p-value from the complex-wide correlation and complex-wide abundance tests on the −log10 scale. CCT-complex is highlighted. (B) Volcano plot for age differences in individual protein abundance for cerebellum with CCT-complex members highlighted with color based on direction of effect. Counts of proteins with significantly higher abundance in older mice and younger mice are included (FDR < 0.1). (C) Pearson correlations from the CCT-complex in younger (left) and older (right) mouse cerebellum. Black and gray squares highlight patterns in the correlation matrix that mirror the structure of the CCT-complex. (D) Histograms of pairwise correlation coefficients between CCT-complex members with each other (top) and other proteins (bottom). Vertical red dashed lines represent median correlations. (E) The CCT-complex is composed of two identical octomeric rings. The CCT2 (*β*) and CCT6A (*ζ*) from each ring are in physical with their twin. Outline of proteins matches correlation structure previously highlighted. (F) GO categories enriched in proteins that are more correlated with CCT-complex members in older mouse cerebellum than younger. The microtubule GO category (GO:0005874) is explored further in Figure S10.

### Ribosomal large subunit complex is more cohesive in young female liver tissue

The CRLS was found to be significantly more cohesive in livers of younger mice (*p* = 4.8e-57) and females (*p* = 2.7e-8) (Figure 7A,C). There was no complex-wide age effect on abundance (*p* = 0.86) or any significant age differences detected for individuals CRLS proteins (FDR < 0.1), whereas female liver had lower complex-wide abundance (*p* = 2.9e-6) and significantly lower abundance for 19 proteins (Figure 7B,D). This replicates our previous finding of decreased ribosomal protein abundance in female liver tissue from genetically diverse mouse populations^15,16^. The co-occurrence of effects on individual protein abundance and complex-wide abundance and cohesiveness led us to examine the age-by-sex interaction effects on individual proteins, where 14 had age-by-sex differences (age-by-sex *p* < 0.05) with a consistent pattern of females having distinctly lower abundance in the younger mouse liver (Figure 7E). These differences contribute to the unique age-by-sex co-abundance patterns of the CRLS in the liver, which is more cohesive in young females compared to older females or males (Figures 7F-G). Notably, age and sex effects on the CRLS vary across the tissues; in lung there is a complex-wide greater abundance in older mice and little clear effect from sex (Figure S11).

**Figure 7.**
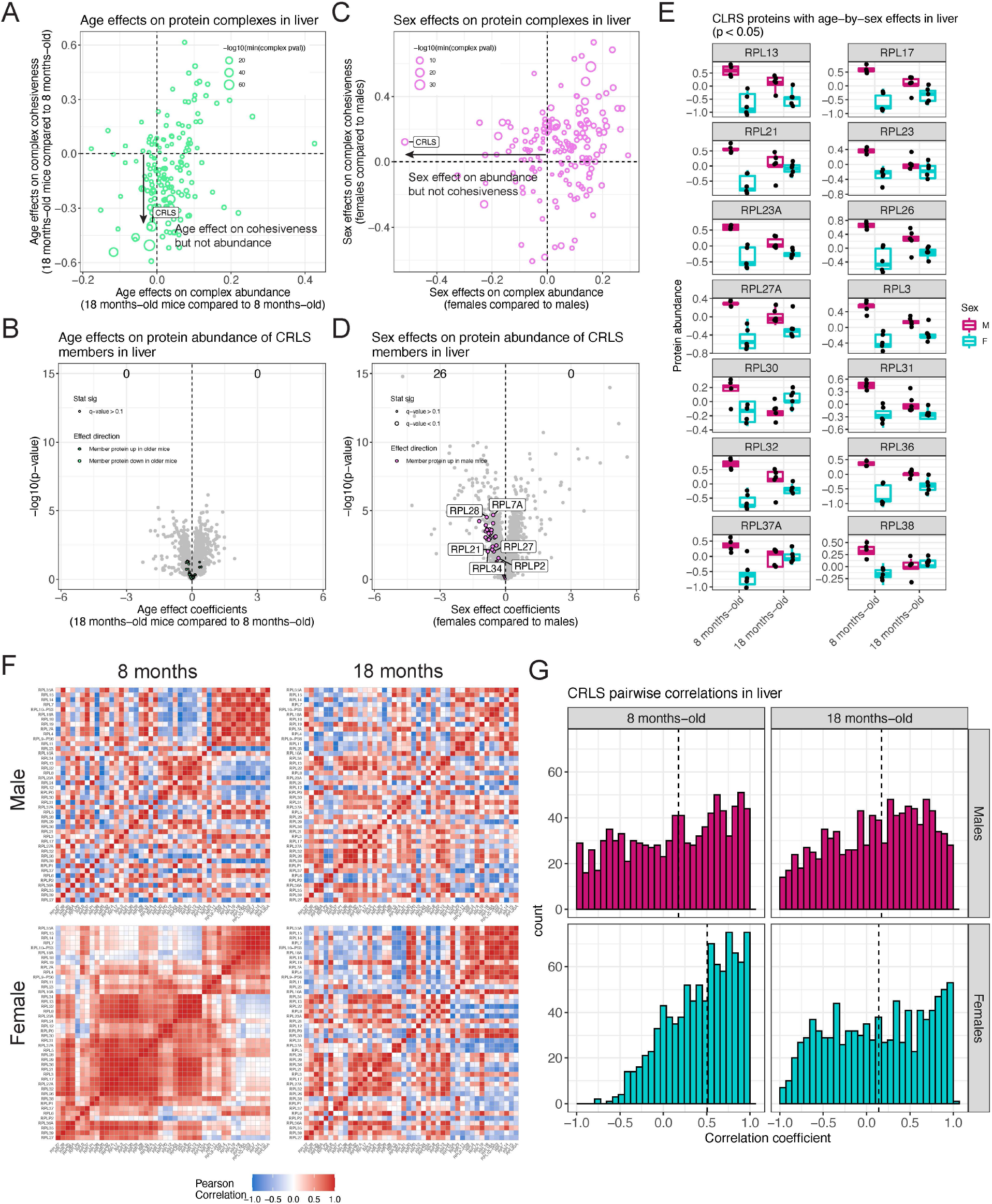
CRLS in liver shows age-by-sex differences in complex-wide abundance and stoichiometry. (A) Comparison of age-related differences in complex-wide correlation (y-axis) to complex-wide abundance (x-axis) in liver. Point size corresponds to the minimum p-value from the complex-wide correlation and complex-wide abundance tests on the −log10 scale. CRLS is highlighted. (B) Volcano plot for age differences in individual protein abundance for liver with CRLS members highlighted with color based on direction of effect. Counts of proteins with significantly higher abundance in older mice and younger mice are included (FDR < 0.1). (C) Comparison of sex-related differences in complex-wide correlation (y-axis) to complex-wide abundance (x-axis) in liver. Point size corresponds to the minimum p-value from the complex-wide correlation and complex-wide abundance tests on the −log10 scale. CRLS is highlighted. (D) Volcano plot for sex differences in individual protein abundance for liver with CRLS members highlighted with color based on direction of effect. Counts of proteins with significantly higher abundance in females and males are included (FDR < 0.1). (E) CRLS proteins with age-by-sex differences in abundance in liver (age-by-sex *p*-value < 0.05). (F) Pearson correlations from the CRLS, stratified by age (younger – left and older – right) and sex (male – top and female – bottom) in mouse liver. (F) Histograms of pairwise correlation coefficients from the CRLS, stratified by age (younger – left and older – right) and sex (male – top and female – bottom) in mouse liver. Vertical dashed lines represent median correlations.

## DISCUSSION

We performed quantitative protein profiling across 10 tissues from the mouse reference strain, C57BL/6J. We tested for and characterized differences in protein abundance based on age, sex, and their interaction. Looking within and across tissues, we identify unique functional patterns of proteins that vary with age, including global changes to components of the immune system, tissue-specific changes to cellular respiration and metabolism, and proteostasis. Furthermore, we examined how protein complexes differ based on age and sex in terms of complex-wide abundance and cohesiveness or correlation.

The primary aging pattern shared across all tissues is an increase in immunoglobulin proteins, implicating the immune system in the aging process. This parallels immune-aging signatures in transcriptomics^2^, and the strongest immune signal for both proteins and transcripts occurs in fat tissues. Even in tissues like striatum and hippocampus that had very few proteins with significant age differences, specific immune proteins increased in abundance with age consistent with the other tissues. Given that the data represent bulk tissue samples, age effects within a tissue may be driven by changes in cellular composition with age. These patterns suggest that with age changes in the adaptive immune response occur, potentially due to increased presence of immunoglobulin-producing immune cell, to varying degrees across tissues. This is further supported by matching increases in the immunoproteasomes.

Additionally, we observed immune-related aging patterns that were specific to tissues, which further highlights changes to the immune system as a signature of aging. In the spleen, which has a unique role in the immune system compared to the other tissues in this study, we saw reduced abundance with age for proteins involved in leukocyte activation, particularly T cells (including ITGB7, SLFN1, SATB1, FOXP1, FOXO1, SIT1, and THY1). We note that many of these proteins were primarily quantified in spleen. Fat also exhibited unique increases with age for immune-related proteins. Together these findings point to dynamic immune system changes across tissues.

Previous studies have demonstrated that protein complexes can be co-regulated by biological factors, such as sex, diet, and genetics^14–16^. We assessed how age and sex affect complex-wide abundance and cohesiveness, replicating our previous finding of reduced ribosomal protein abundance in female liver^15^. We note an excess of age differences across tissues for protein complexes, both in terms of cohesiveness and abundance, related to proteostasis, which is hypothesized to contribute to aging^40–43^, including ribosomes, proteasome, and the CCT-complex. We note that the direction of change was not necessarily consistent across tissues for a complex, nor is it possible to distinguish loss of stoichiometric balance with age from changes in tissue compositions, as is likely the case for the immunoproteasome. Nevertheless, our findings reveal changes to and potential disruption of proteostasis with age that vary across tissues.

Further studies are needed to understand the mechanisms underlying the large-scale aging dynamics of proteins discovered here, many of which are independent of transcription. A broader time series of age groups, similar to the gene expression study of Schaum *et al* 2020^2^ would provide a higher resolution picture of how protein abundance changes with age. Such a study could characterize non-linear trends of aging, which could then be compared between groups of age co-regulated proteins or even to aging trends in gene transcripts.

Without corresponding gene expression data from the same samples, it is experimentally challenging to distinguish whether aging changes in proteins observed in bulk tissue reflect consistent changes within cells or changes in tissue composition. If gene expression data were available, tissue deconvolution^45^ of bulk tissue RNA-seq would be possible using single cell data from overlapping tissues from a resource, such as the *Tabula Muris*^3^, to estimate cell-type proportions per sample. The relationship between cell-type proportions and age could then be tested to identify proteins with age effects that correspond to changes in tissue composition.

Even in the absence of gene expression data, there are hints that some aging effects on proteins reflect changes to tissue composition with age. For example, increased levels of immunoproteasomes and decreased levels of complement cascade proteins across multiple tissues could be explained by a changing balance of immune cells with age. Estimation of cell-type identity through integration of reference single cell RNA-seq with bulk protein abundance rather than gene expression is challenging because fewer genes are measured at the protein level, resulting in likely less information distinguishing cell-types. Furthermore, single cell gene expression and mass-spec proteomics are less comparable. Single cell proteomics data for samples could more directly elucidate age changes in proteins at a cellular level. However, these approaches are newly developing^46–48^ and pose both technical and analytical challenges to overcome, such as extreme data sparsity.

Based on our prior studies, we sought to evaluate the effect of aging on proteins across a wide range of tissues in the reference mouse strain to assess the concordance of age effects on proteins and their transcripts across tissues, as well as identify global and tissue-specific patterns of aging at the protein level. This study functions as a unique quantitative protein resource for the aging-focused research community. We provide our data and corresponding processed results as an interactive RShiny application, available online at http://aging-b6-proteomics.jax.org, allowing proteins of interest to be easily queried. This tool can be used to confirm or replicate findings from previous studies in mice (BCAT1) or other models (VCAM1 in humans and REN1 in aging rat kidney) and generate hypotheses for future studies of aging.

## METHODS

### Data and code availability

The mass-spec proteomics data for all samples reported here have been deposited in ProteomeXchange (http://www.proteomexchange.org/) via the PRIDE partner repository (ProteomeXchange: PXD034029). All statistical analyses were performed using the R statistical programming language (v4.0.3)^49^. All starting data, key forms of processed data, and the analysis pipeline to process the data, run analysis, and produce the reported findings have been made publicly available (https://doi.org/10.6084/m9.figshare.19765849).

The processed data are also interactively viewable through an RShiny application, which is available online (http://aging-b6-proteomics.jax.org) or can be run locally through RStudio (https://github.com/gkeele/Aging_B6_Proteomics_RShiny).

### Mice

Female and male C57BL/6J mice (stock JR#000664) were obtained from The Jackson Laboratory. Animals were maintained on pine shavings and given a standard rodent diet (LabDiet 5KOG) and acidified water in a pathogen free room. The room was maintained at 21°C with a 12-hour light/dark cycle (6am to 6pm). At the time of tissue collection (at 8 and 18 months of age) animals were euthanized by cervical dislocation. Kidney, liver, fat (inguinal adipose), spleen, lung, heart, skeletal muscle (quadriceps), striatum, cerebellum, and hippocampus tissues were collected from each animal. All animal experiments were performed in accordance with the National Institutes of Health Guide for the Care and Use of Laboratory Animals (National Research Council) and were approved by The Jackson Laboratory’s Animal Care and Use Committee.

### Sample preparation for proteomics analysis

Tissue samples were dounce homogenized and resuspended in lysis buffer (8M urea, 150 mM NaCl, Roche protease inhibitor tablets) and cells were lysed by sonication (procedure). Lysates were cleared by centrifugation (15 min at 20,000×g) and protein concentrations were measured using Pierce BCA assay kits. Proteins were then reduced with dithiothreitol (5mM for 30 minutes at room temperature) and alkylated with iodoacetamide (15mM for 60 minutes in the dark). The alkylation reaction was quenched by adding an additional aliquot of DTT. For each sample, 100ug of protein was aliquoted and diluted to a final concentration of 1mg/mL. Proteins were digested using LysC (Wako, overnight, room temperature, moderate agitation) followed by trypsin (6 hr, 37C, 200rpm). The resulting peptides were then labeled with individual TMT (Thermo) reporters (1.5 hours at room temperature) and the reaction was quenched with hydroxylamine (5% in water for 5 minutes). Labeled peptides were mixed into a set of two plexed for each tissue analysis. After labeling and mixing, peptide mixtures were desalted using C18 seppak cartridges (1mg, Waters). Desalted peptides were then fractionated using basic-pH reverse phase chromatography^50^. Briefly, peptides were resuspended in Buffer A (10mM ammonium bicarbonate, 5% acetonitrile [ACN], pH 8) and separated on a linear gradient from 13% to 42% Buffer B (10mM ammonium bicarbonate, 90% acetonitrile [ACN], pH 8) over an Agilent 300Extend C18 column using an Agilent 1260 HPLC equipped with single wavelength detection at 220nm). Fractionated peptides were desalted using Stage-tips^50^ prior to LC-MS/MS analysis.

### Mass spectra data analysis

Peptides were separated prior to MS/MS analysis using an Easy-nLC (Thermo) equipped with an in-house pulled fused silica capillary column with integrated emitter packed with Accucore C18 media (Thermo). Separation was carried out with 90-minute gradients from 96% Buffer A (5% ACN, 0.125% formic acid) to 30% Buffer B (90% ACN, 0.125% formic acid). Mass spectrometric analysis was carried out on an Orbitrap Fusion Lumos (Thermo). Multiplexed analysis of samples was done using real-time search data acquisition^10^, based on canonical SPS-MS3 acquisition. Briefly, a survey MS1 scan was used to identify potential peptide precursors (R = 120000, Mass range: 400-2000 m/z, max Inject time: 50ms, AGC: 200000, RF lens: 30%). The top 10 precursors were selected for fragmentation and analysis in the ion trap (Dynamic exclusion: 120s at 10ppm, CID collision energy: 35%, max inject time: 120ms, AGC: 20000, scan rate: rapid, isolation width: 0.5 m/z). Real-time spectral matching was carried out using the Comet search algorithm^51^. If, and only if, a peptide was matched with high confidence, the instrument would then acquire an additional SPS-MS3 scan for quantification of relative abundances (R = 50000, HCD NCE: 65, max inject time: 200ms).

Raw spectral information was converted to mzXML format using Monocle^52^, and spectra were matched using the Comet search algorithm comparing against the ENSEMBL_GRCm39 database^51,53^. Peptides and proteins were filtered to a 1% using rules of protein parsimony^51^.

### Protein abundance estimation from peptides

Samples from each tissue were run across two tissue-specific batches. For each tissue, the abundance level for proteins was estimated as a scaled sum of their component peptides. For protein *j* from tissue *k* of mouse *i*, the abundance is calculated as 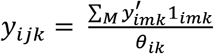 where *M* is the set of peptides that map to protein 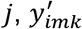 is the intensity of peptide *m* from tissue *k* of mouse *i*, 1*i_mk_* is the indicator function that peptide *m* was observed in tissue *k* of mouse *i*, and 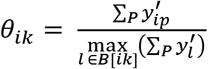 is the within-batch scaling factor^54^ for mouse *i* in tissue *k*, representing the ratio of the sum of all peptide intensities for mouse *i* to the maximum sum total for the 11 samples in the batch of mouse *i* for tissue *k*, *B*[*ik*]. To standardize quantities across the two batches, abundances were ratio normalized to a pooled sample that was included in both batches: 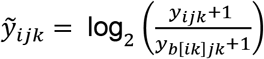 where *b*[*ik*] is the bridge sample from the batch of mouse *i* for tissue *k*.

### Filtering out low quality proteins and samples

We filtered out proteins that were only observed in one of the two batches for a tissue because we found single batch proteins could be influential in downstream analysis. After protein abundance estimation and removal of single batch proteins, we performed PCA^55^ to identify tissue samples that were clear outliers across many proteins. We removed one sample from liver (young male), fat (young male), spleen (old male), lung (young male), and skeletal muscle (old female). Two samples were removed from striatum, both old female mice.

### Testing for age, sex, and age-by-sex interaction effects on proteins within tissues

We tested for significant differences based on age, sex, and age-by-sex using ordinary least squares (OLS) regression. For each protein *j* observed in tissue *k*, we fit the following model:

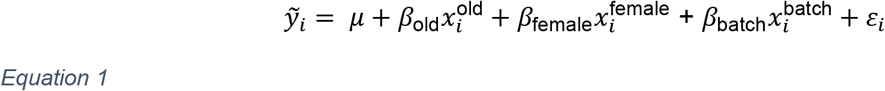

where *μ* is the intercept, *β*_old_ represents the effect of being in the old group, *β*_female_ represents the effect of being female, *β*_batch_ represents the effect of being in second batch, 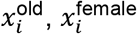, and 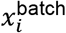 are indicator variables that mouse *i* is old, female, or in batch2, respectively, and *ε_i_* is the error for mouse *i*, modeled as *ε_i_* ~ *N*(0, *σ*^2^) and *σ*^2^ is the variance of the noise. To test for an age effect, we used analysis of variance (ANOVA), comparing the model in Equation 1 to a model excluding the age term and recorded the age effect coefficients, standard errors, and *p*-values. Similarly, we tested for a sex effect by comparing the Equation 1 model to a model excluding the sex term and again recorded sex effect coefficients, standard errors, and *p*-values. Finally, we assessed age-by-sex differences by adding age-by-sex interaction term to the model which was then compared to the Equation 1 model with ANOVA and recorded the age-by-sex *p*-value. We estimated the FDR using the Benjamini-Hochberg (BH) method^56^ for each effect type, producing age, sex, and age-by-sex *q*-values. This process was used each of the 10 tissues.

When plotting data (not effect parameters), we first regressed out the effect of batch to make the signal from age or sex clearer. We fit the model in Equation 1 and then calculated 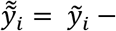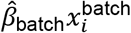, where 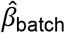 is the estimated coefficient for the second batch, for all proteins *j* across all tissues *k*.

### Testing for consistent and unique age and sex effects on proteins across tissues

We tested whether age and sex effects on a protein were similar or distinct across tissues for all proteins detected in more than one tissue using linear mixed effects models (LMM) fit in the lme4 R package^57^. For each protein *j*, we fit the following model:

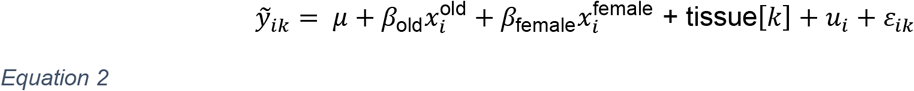

where tissue[*k*] represents the effect of tissue *k*, *u_i_* is a random term specific to mouse *i*, modeled as *u_i_* ~ *N*(0, *τ*^2^), *τ*^2^ is the variance component underlying the mouse-specific effect, and all other terms as previously defined. A batch effect was not included because tissue and batch are highly confounded (batch pairs specific to each tissue), and we are not interested in the marginal effect of either tissue or batch. We tested for a consistent age effect across tissues by comparing the model in Equation 2 to reduced model excluding the age effect with ANOVA using Satterthwaite’s approximation for an LMM^58,59^, implemented in the lmerTest R package^60^. We next looked for age effects that were unique to tissues or were even flipped by testing an age-by-tissue term by comparing the model in Equation 2 to an expanded model that included the interaction term. The same approach was used to test for consistent sex effects across tissues and tissue-unique and flipped sex effects. To account for multiple testing across proteins, we again estimated *q*-values using the BH method^56^.

### Modeling how factors contribute to variation in proteins measured in all 10 tissues

To evaluate how various technical and biological factors contribute to variation in the abundance of individual proteins, we fit an LMM with variance components for each factor for all proteins measured in all 10 tissues. For each protein *j*, we fit the following model:

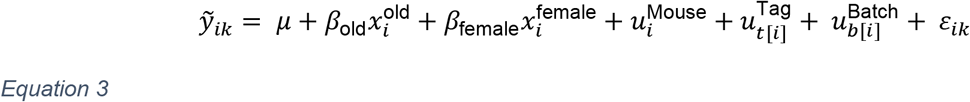

where 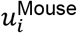 is a random term specific to mouse *i* (20 levels), 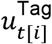 is a random term for TMT tag *t* of mouse *i* (10 levels), 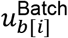 is a random term for TMT batch *b* of mouse *i* (20 levels), and all other terms as previously defined. Each random effect is modeled as 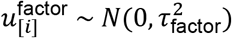. Proportion of variation explained by each factor was calculated as 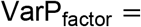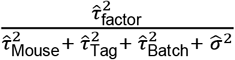. For point and interval estimates of the random terms, best linear unbiased predictors (BLUPs) and 95% predictive intervals were used.

### Testing for age and sex effects on transcripts within tissues

We obtained transcriptomics data reported in Schaum *et al* 2020^2^, which represent 17 tissues and 10 age groups (1 to 27 months) from the closely related C57BL/6JN (B6N) mice. Each age group consisted of samples from four males and two females, except for 24 and 27 months, which only had the four males. We filtered the data to the 9 and 18 months-old age groups, which most closely match our age groups of 8 and 18 months. We then tested for and characterized age and sex effects (log fold change) on gene expression within each tissue. We used the DESeq2 R package^61^ to fit models similar to Equation 1 (excluding the batch covariate), but now using a negative binomial generalized linear model (GLM) to accommodate that the data are gene counts. Age and sex effects on proteins and transcripts were compared based on aligning Ensembl gene IDs. When plotting the data to illustrate effects, we first used DESeq2’s variance stabilizing transformation^62^ across samples from all tissues and age groups.

### Summarizing protein complex co-abundance

We quantified how co-abundant, *i.e*., cohesive, a protein was with its complex^36^ as the median Pearson correlation coefficient between it and other complex members. An overall summary for the complex was then taken as the median across all the individual medians, an approach we used previously^15^. We calculated cohesiveness only for protein complexes with more than three member proteins observed for a given tissue.

### Testing for consistent age and sex effects on abundance across a protein complex

We tested for consistent age and sex effects on protein complexes with more than three observed members by jointly modeling all proteins. For each protein complex *c* observed in tissue *k*, we fit the following LMM for proteins *j* ∈ *J_c_*:

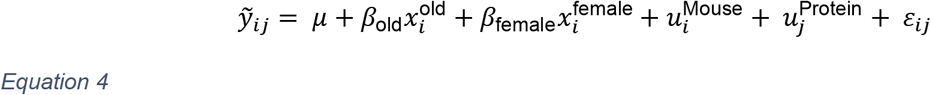

where *J_c_* is the set of proteins in complex *c*, 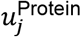 is a random term for protein *j*, modeled as 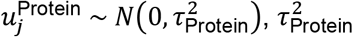 is the variance component underlying proteins, and all other terms as previously defined. We tested for an age or sex effect by comparing the model in Equation 4 to either a model excluding the age or sex term, respectively, through ANOVA, again with Satterthwaite’s approximation^58,59^, producing a *p*-value. FDR was again estimated using the BH method^56^ across protein complexes and tissues, for age and sex separately.

### Testing for age and sex effect on protein complex cohesiveness

Age or sex could affect how tightly co-abundant a protein complex is. We evaluated whether age or sex had a consistent effect on the correlation structure of the complex using a paired *t*-test. For example, with age, we calculated all the pairwise correlations between members for both age groups for a given complex within a tissue: (***r***_Old_, ***r***_Young_), resulting in a *t*-test *p*-value and effect. This process was repeated based on sex. The BH method^56^ was again used to estimate FDR and produce *t*-test *q*-values for both age and sex. We used a more stringent significance threshold of FDR < 0.01 because correlation coefficients are non-standard quantities to model with a *t*-test.

### Testing changes in correlation between pairs of proteins with permutations

To test whether individual pairwise correlations between protein complex members differed based on age or sex, we used permutations. When testing for an age difference, we swapped mouse identifiers among males and among females, thus maintaining the effect of sex and potentially avoiding anti-conservative permutation *p*-values. When testing for a sex difference, we swapped labels while maintaining the age groups. We estimated empirical *p*-values as 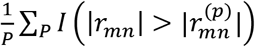 where *P* is the number of permutations, *I*(.) is an indicator function, |*r_mn_* | is the observed absolute Pearson correlation coefficient between proteins *m* and *n*, and 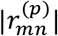 is the absolute Pearson correlation coefficient for permutation *p*. We set *P* to 100,000.

### Gene set enrichment analyses

We used both set-based and score-based gene set enrichment analyses and confirmed that they often conferred with each other. We used the clusterProfiler R package^12^ for set-based analysis, in which we defined gene sets based on various criteria, such as all genes from a given tissue that have significant age or sex effects on protein abundance compared to all genes analyzed in the tissue. We also evaluated tissue-specific gene sets defined by the direction of the significant age or sex effect for given tissues or based on how genes clustered according to age effects across the tissues. For the score-based analysis, we used the fgsea R package^63^ paired with GO pathways from the msigdbr R package^64^. For each tissue, we input all analyzed genes with scores as the age or sex coefficients from Equation 1, standardized by their standard errors. We compared GO findings between pairs of tissues by intersecting tissue-specific results based on pathway ID and gene ID.

## ACKNOWLEDGEMENTS

We gratefully acknowledge Vivek M. Philip and the Computational Sciences Service at The Jackson Laboratory for assistance in the development of the interactive RShiny tool for querying the data. This work was supported by grant funding from the National Institutes of Health (NIH): F32GM134599 (G.R.K.), R01GM067945 (S.P.G.), and P30AG038070 from The Jackson Laboratory Nathan Shock Center of Excellence in the Basic Biology of Aging (R.K. and G.A.C.).

## AUTHOR CONTRIBUTIONS

R.K., S.P.G., G.A.C., and D.K.S. conceptualized the project and designed the experiment. J.S. and D.K.S. performed the experiments. G.R.K and G.A.C. designed the methodology. G.R.K. performed the analysis. G.R.K. and J.Z. built the interactive webtool. G.R.K., R.K., G.A.C., and D.K.S. wrote the manuscript. All authors reviewed the manuscript.

## Notes

https://figshare.com/articles/dataset/Data_and_code_for_aging_C57_B6J_proteomics_manuscript/19765849

https://github.com/gkeele/Aging_B6_Proteomics_RShiny

### Competing Interest Statement

The authors have declared no competing interest.

### Summary of Updates

We revised the manuscript to improve clarity. More material on potential future experiments and limitations have been added to the Discussion.

