## Supplemental Figures 1-11 for "Global and tissue-specific aging effects on murine proteomes"

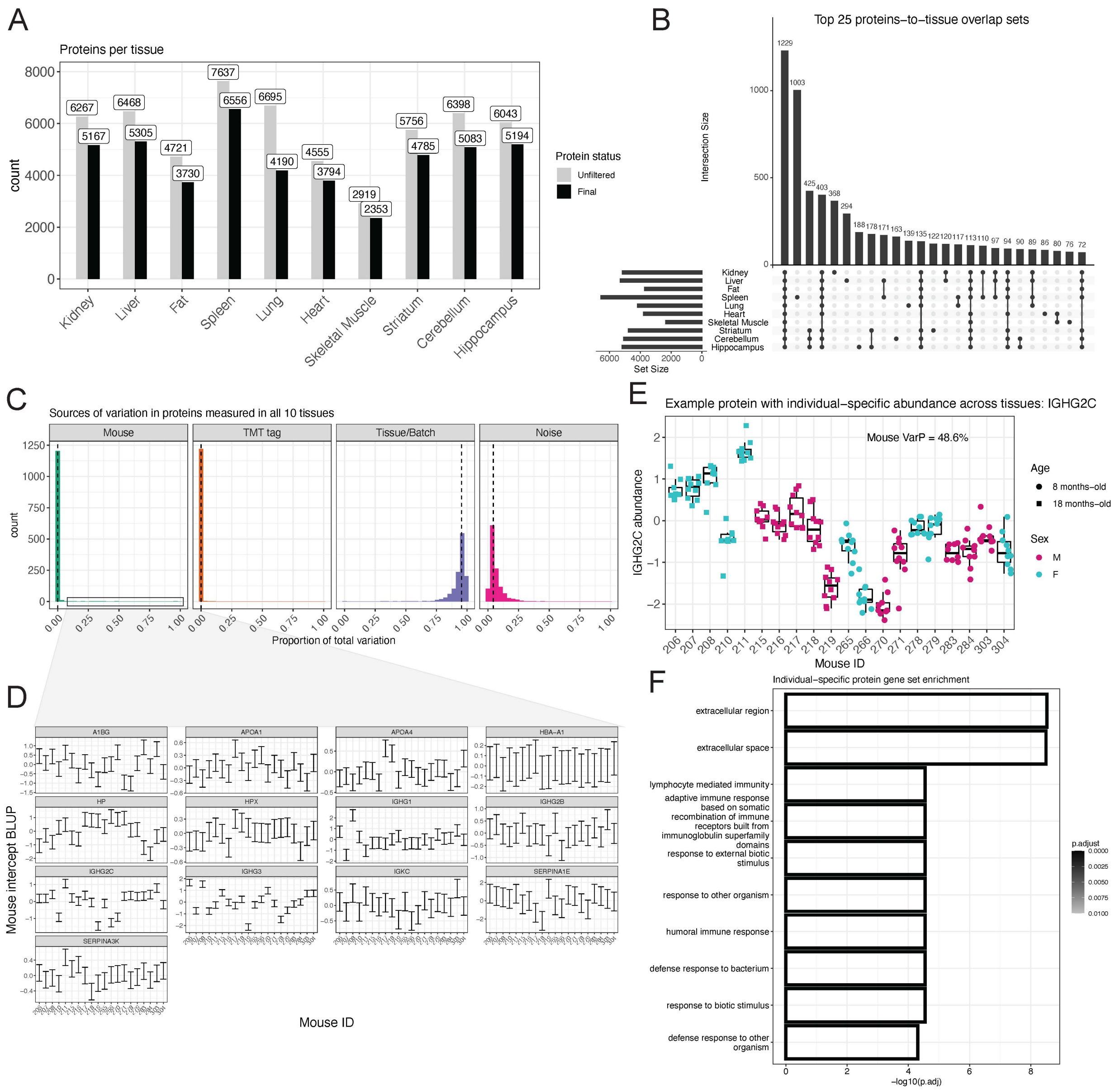


**Figure S1. Proteins with individual-specific abundance across 10 tissues.**
(A) The total number of proteins quantified and analyzed after filtering out proteins detected in only a single batch for a given tissue.
(B) The top 25 intersections of proteins and the tissues in which they were detected and analyzed.
(C) Histograms of the proportion of variance explained by mouse, TMT tag, tissue/batch, and noise for the 1,229 proteins analyzed in all 10 tissues. Vertical dashed lines represent median proportions. Tissue and batch are confounded because all 20 samples for a tissue were measured across two tissue-specific batches. The gray square highlights proteins for which a large proportion of variation in abundance (5-95%) is explained by individual mouse ID.
(D) Modeled individual-specific abundance levels for all 20 mice, estimated as 95% prediction intervals of the best linear unbiased predictors (BLUP), for the 13 proteins that mouse ID identity explained 5-95% of abundance variation.
(E) The immunoglobin IGHG2C had highly individual-specific abundance across the 10 tissues. Data are colored by sex and point shape corresponds to age.

(F) GO categories enriched in the 13 individual-specific proteins.

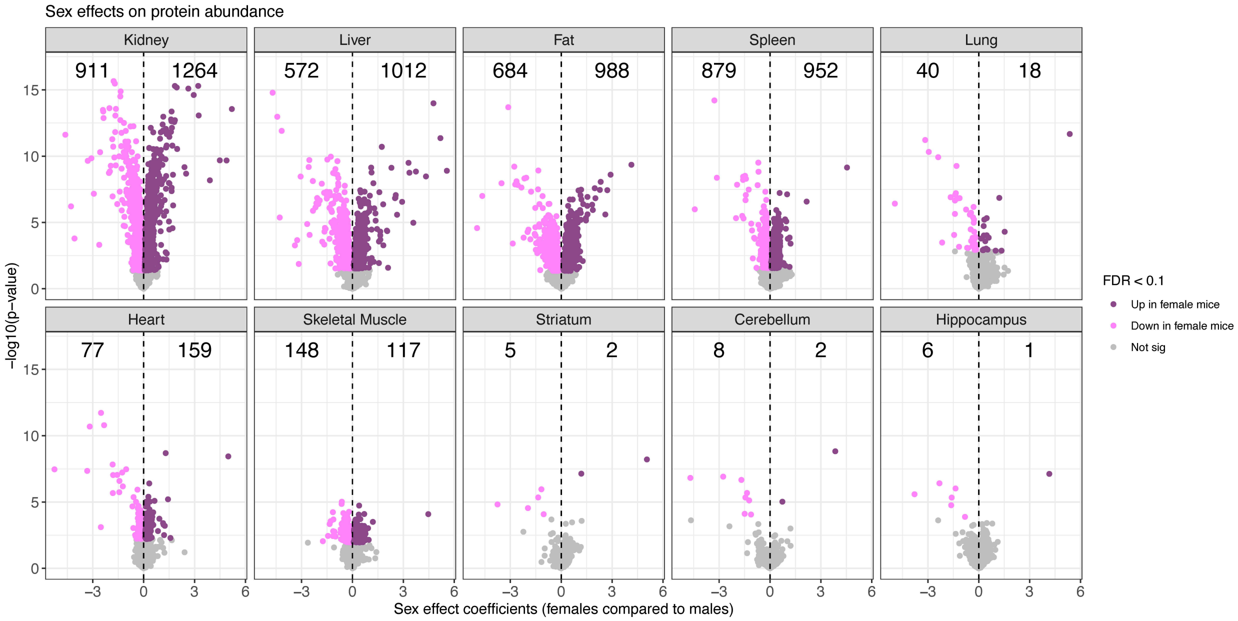


**Figure S2. Proteins with sex-related differences in abundance, across 10 tissues.** Proteins with sex differences in abundance, represented as volcano plots. Differences in protein abundance are summarized as regression coefficients (x-axis) and corresponding -$\log_{10}$(*p*-value) (y-axis). Points are colored based on statistical significance (FDR < 0.1) and direction of effect. Counts of proteins with significantly higher abundance in females and males are included. Dashed vertical lines at 0 included for reference. Proteins with abundance age differences are shown in Figure 2.

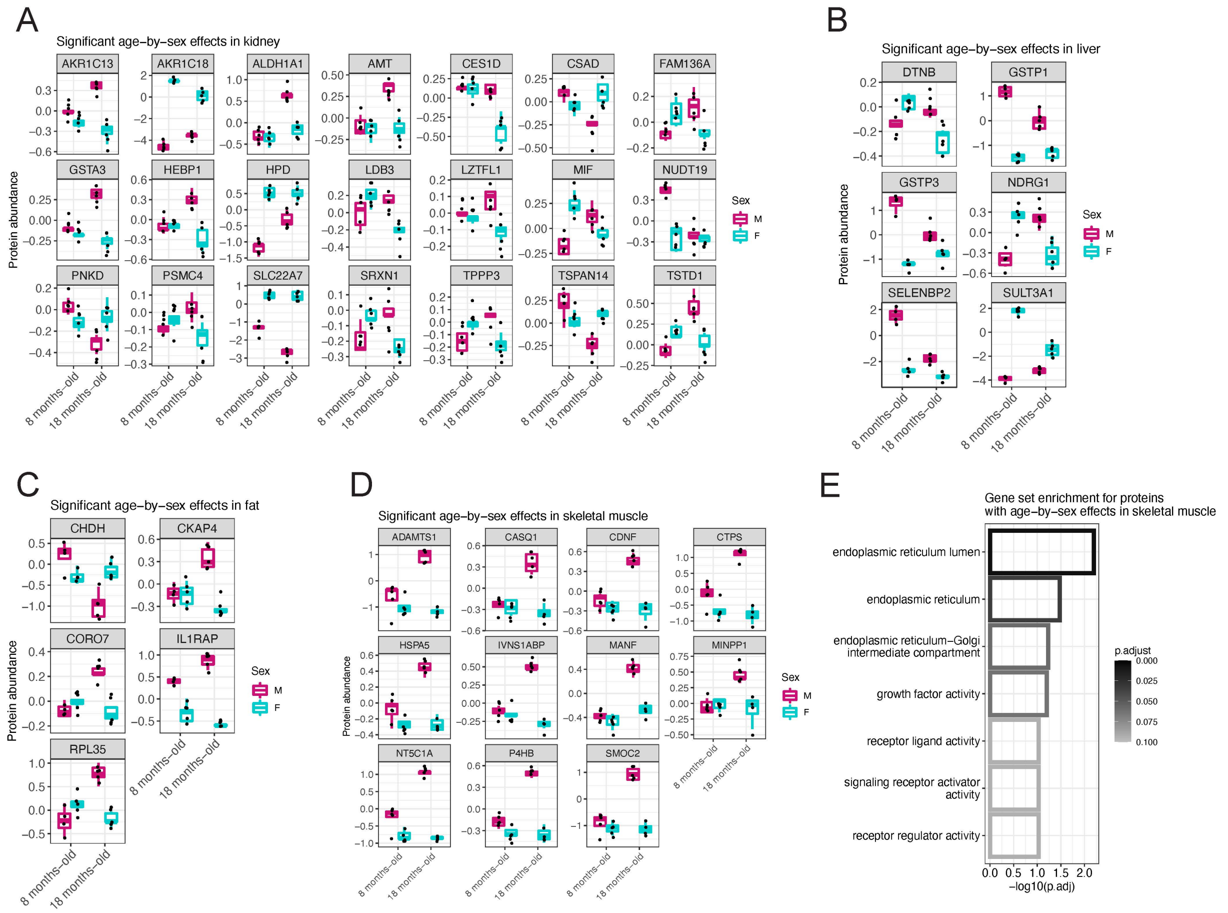


**Figure S3. Proteins with age-by-sex interaction effects in abundance (FDR < 0.1).** Proteins with sex-by-age interaction effects for (A) kidney, (B) liver, (C) fat, and (D) skeletal muscle. (E) Proteins with statistically significant GO categories enriched in proteins with age-by-sex interaction effects in skeletal muscle.

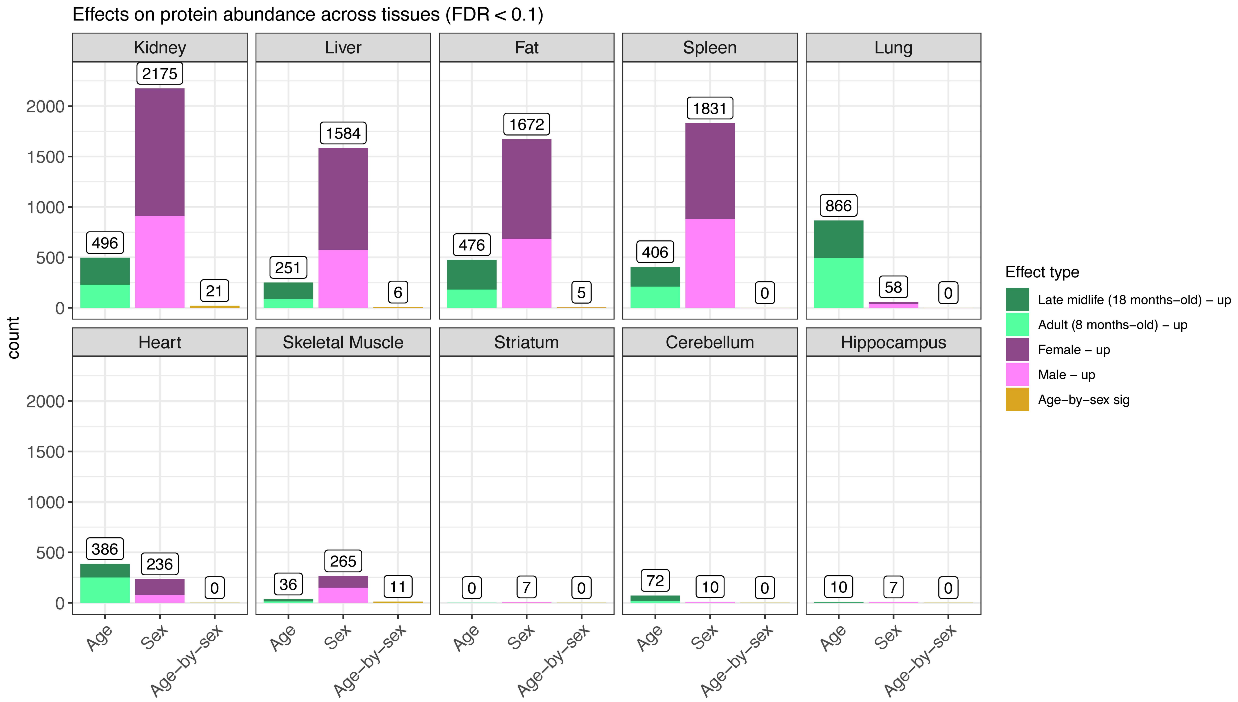


**Figure S4. Counts of proteins with age, sex, and age-by-sex differences (FDR < 0.1), across 10 tissues.**

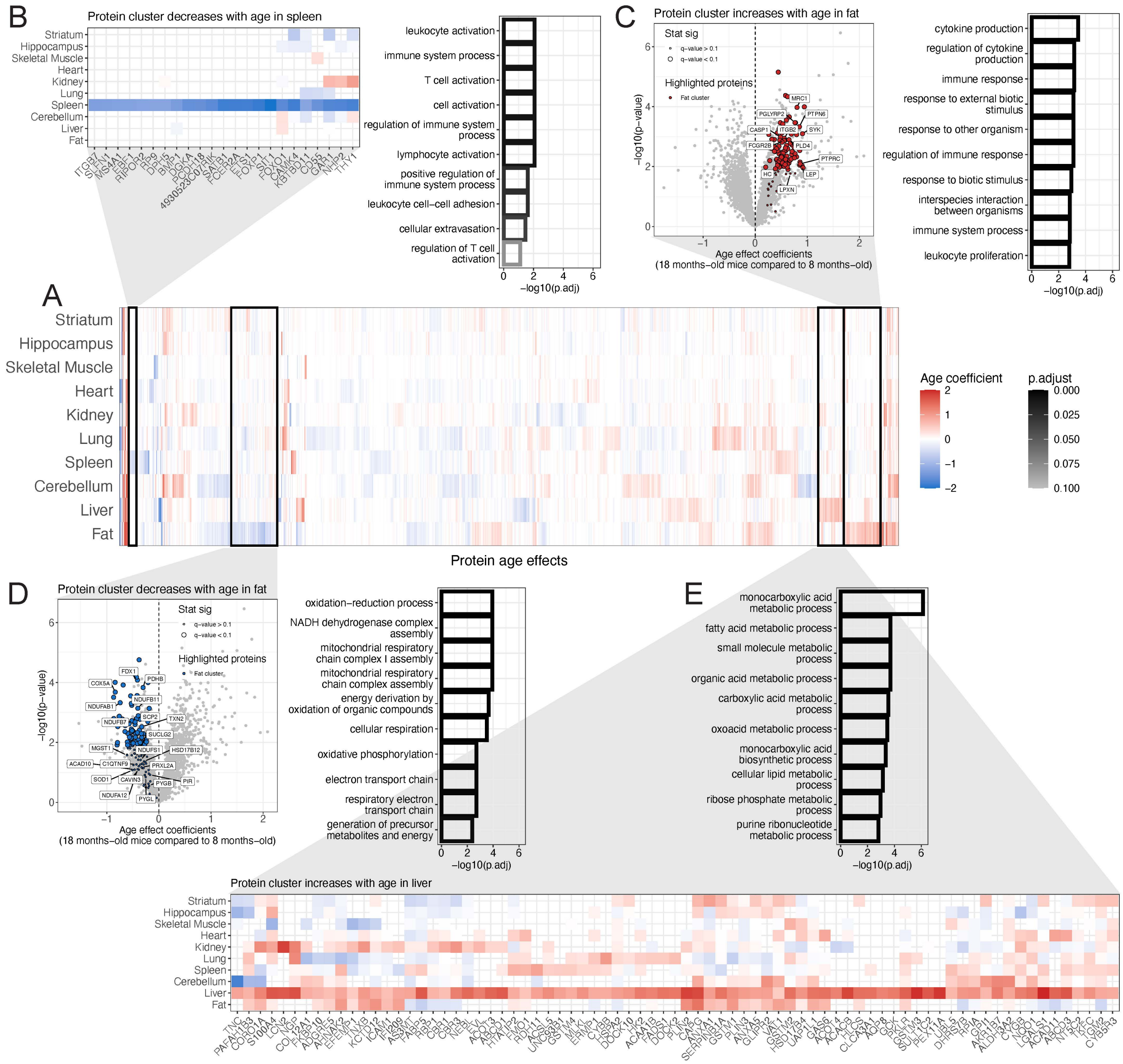


**Figure S5. Additional patterns of aging unique to specific tissues.**

(A) Age differences detected across the 10 tissues (FDR < 0.1), represented as a heatmap. Differences are summarized as regression coefficients. Hierarchical clustering of the proteins (columns) reveals sets of proteins with age difference patterns that are unique to specific tissues.

Zoomed-in heatmaps are shown for (B) spleen-specific decreased abundance in proteins related to immune activation in older mice and (E) liver-specific increased abundance in proteins related to a number of metabolic processes in older mice. Volcano plots are shown for (C) fat-specific increased abundance in proteins related to immune response and cytokine production in older mice and (D) fat-specific decreased abundance in proteins related to cellular respiration and the mitochondrial respiratory chain complex in older mice. Gene set enrichments are shown for all tissue-specific gene sets.

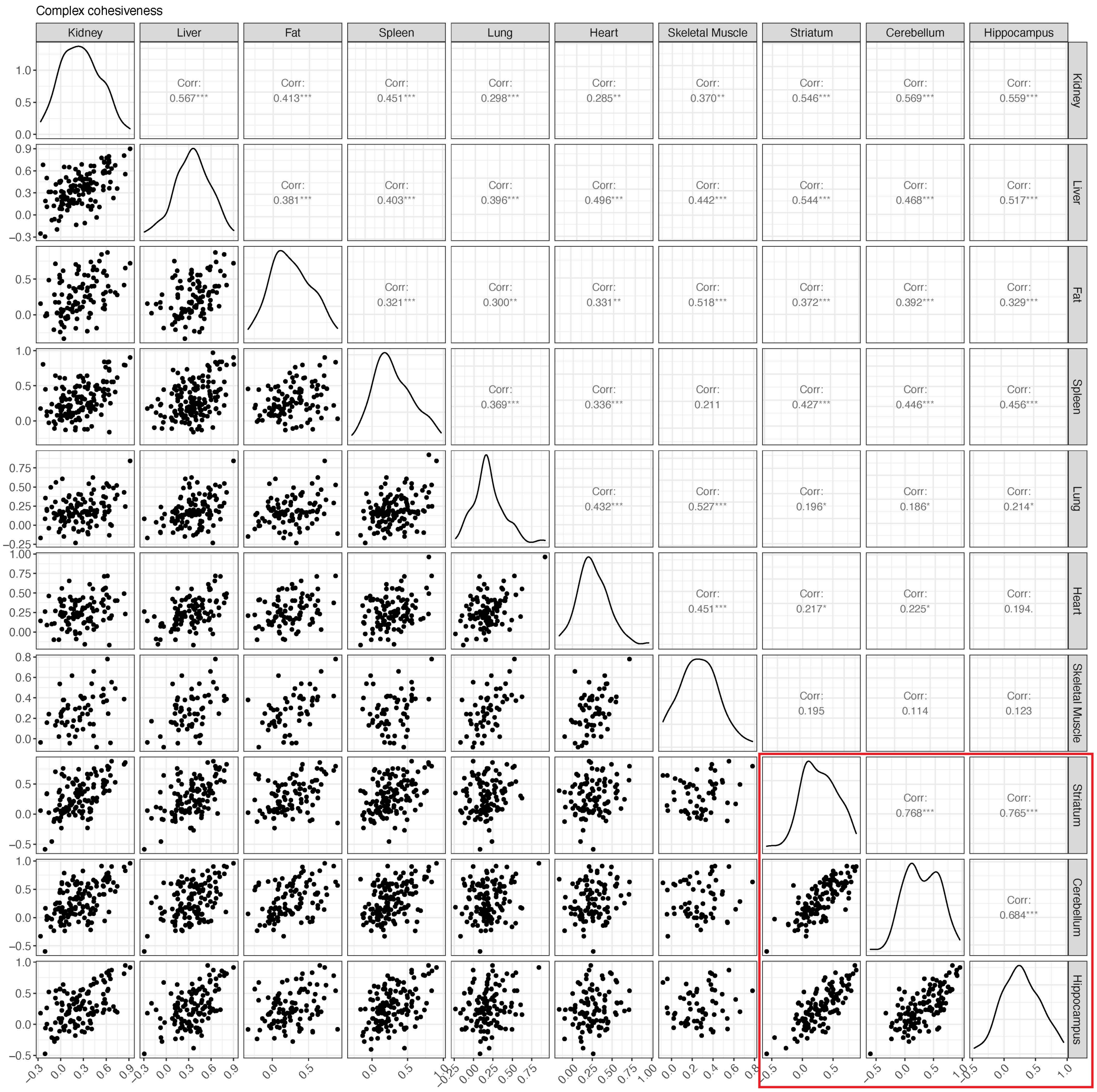


**Figure S6. Comparison of protein complex cohesiveness between tissues.** Cohesiveness for protein complexes is summarized as the overall median across all median pairwise correlations for individual proteins with other complex members. The red square highlights the three brain tissues.

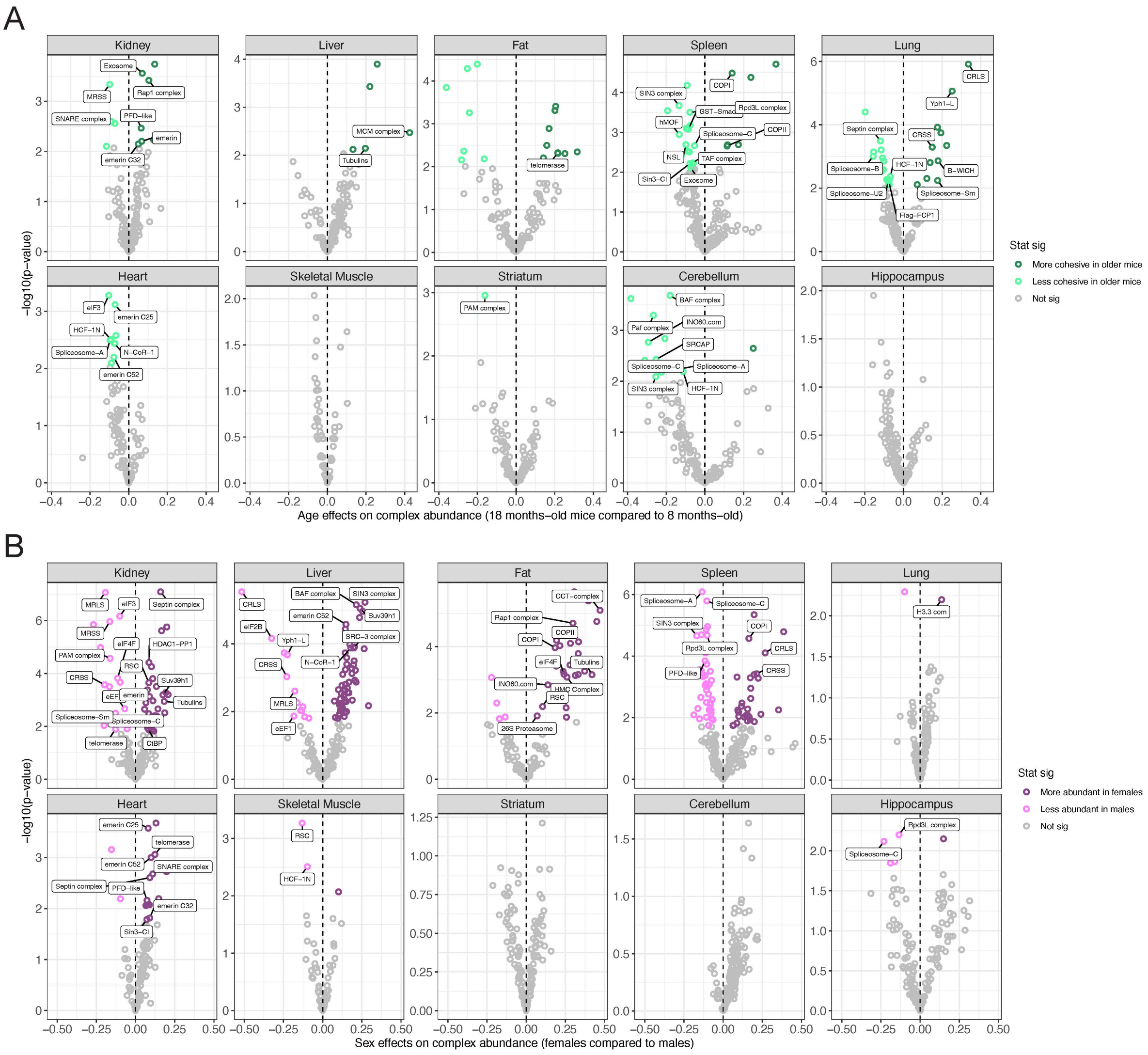


**Figure S7. Protein complexes with age- and sex-related differences in complex-wide abundance, across 10 tissues.** Protein complexes with (A) age-related and (B) sex-related differences in complex-wide abundance, represented as volcano plots. Differences in protein abundance are summarized as regression coefficients (x-axis) and corresponding -$\log_{10}$(*p*-value) (y-axis). Points are colored based on statistical significance (FDR < 0.1) and direction of effect. Dashed vertical lines at 0 included for reference.

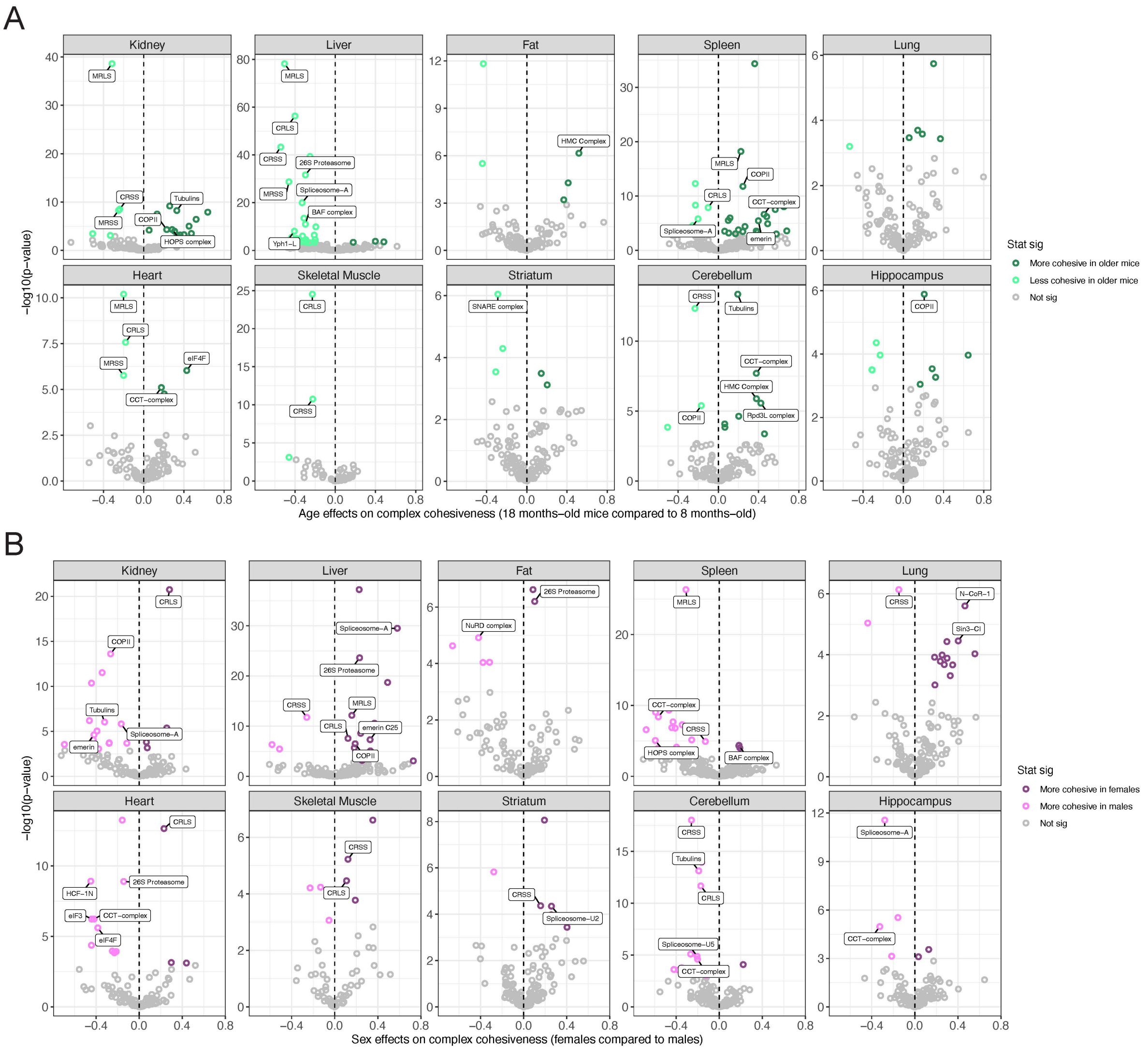


**Figure S8. Protein complexes with age- and sex-related differences in stoichiometry, across 10 tissues.** Protein complexes with (A) age-related and (B) sex-related differences in complex-wide correlation, represented as volcano plots. Differences in complex-wide correlations are summarized as the effect estimate from a paired *t*-test (x-axis) and corresponding -$\log_{10}$(*p*-value) (y-axis). Points are colored based on statistical significance (FDR $< 0.01$) and direction of effect. Dashed vertical lines at 0 included for reference.

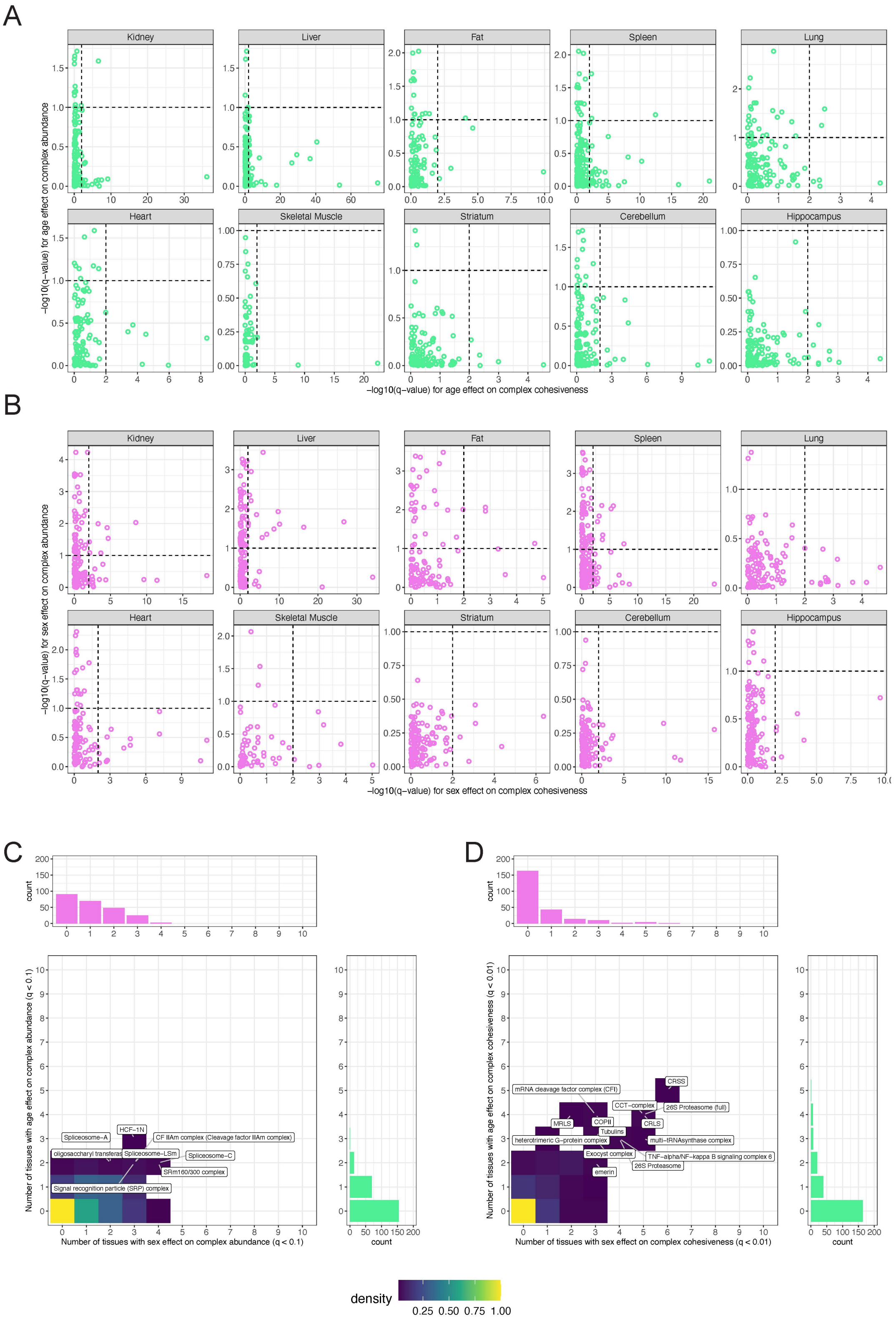


**Figure S9. Comparison of protein complex age- and sex-related differences in complex-wide abundance and stoichiometry, across 10 tissues.**

Comparisons of -$\log_{10}$(*q*-values) for complex-wide abundance differences by complex-wide correlation differences due to (A) age and (B) sex. Vertical dashed line is included at 1 to denote significance for complex-wide abundance differences. Horizontal dashed line is included at 2 to denote significance for complex-wide correlation differences.

Comparison of the number of tissues that had a significant difference due to age (y-axis) and sex (x-axis), for (C) complex-wide abundance and (D) complex-wide correlation. Protein complex with significant differences in 2 or more tissues for both age and sex are labeled.

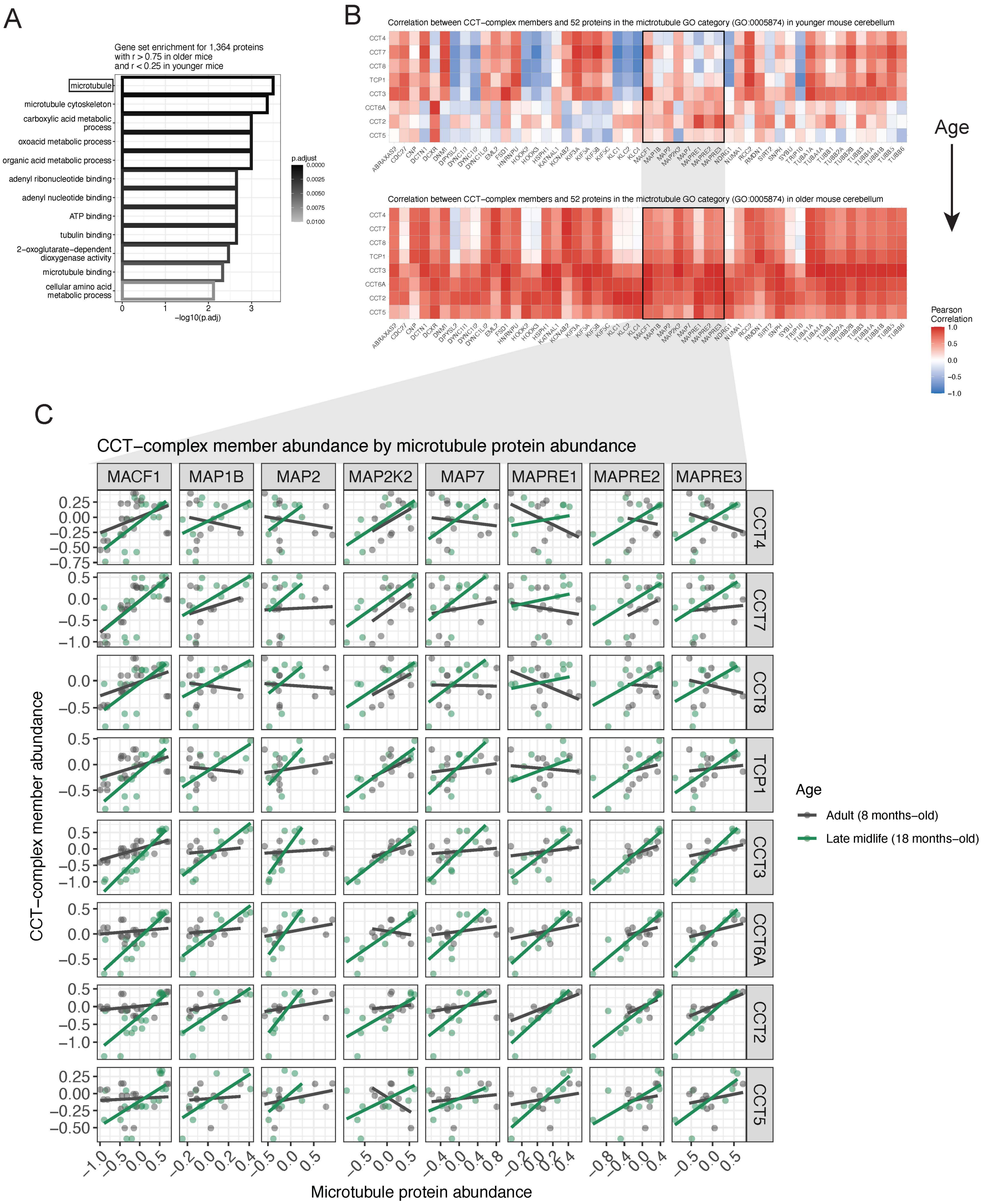


**Figure S10. Microtubule proteins are more correlated with chaperonin-containing T-complex (CCT-complex) members in older mouse cerebellum.**
(A) GO categories enriched in proteins that are more correlated with CCT-complex members in older mouse cerebellum than younger. The microtubule GO category (GO:0005874) is highlighted.

(B) Pearson correlations between CCT-complex members and 52 microtubule proteins (GO:0005874) in younger (top) and older (bottom) mouse cerebellum. Black squares highlight eight microtubule-actin proteins.
(C) Protein abundance of the eight CCT-complex members compared to the eight microtubule-actin proteins. Best fit lines included to reflect correlations. Points and lines are colored based on age.

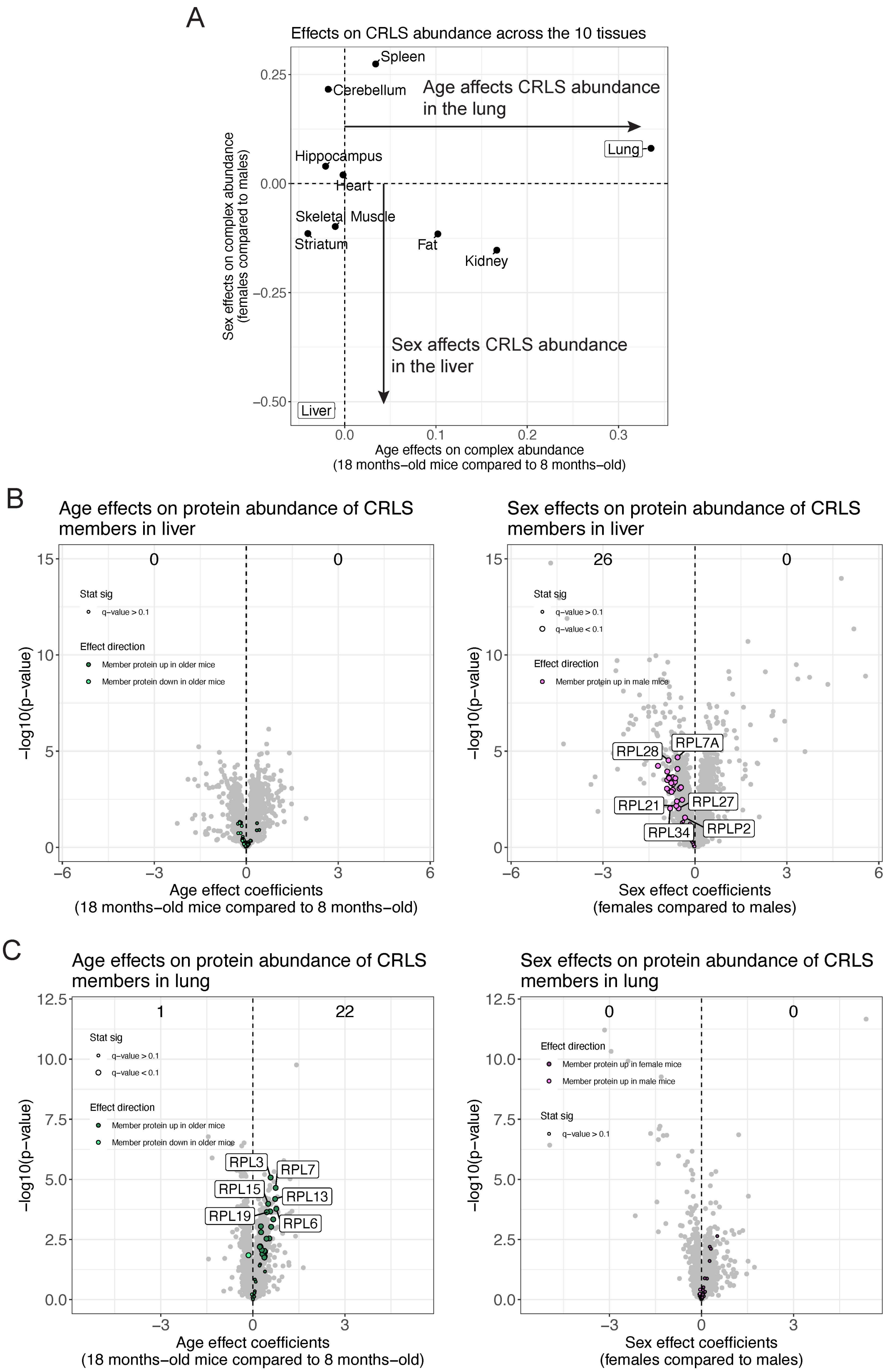
A
Effects on CRLS abundance across the 10 tissues
Spleen
Cerebellum
Age affects CRLS abundance in the lung
Lung
Hippocampus
Heart
Skeletal Muscle
Striatum
Fat
Kidney
Sex affects CRLS abundance in the liver
Liver
Sex effects on complex abundance (females compared to males)
Age effects on complex abundance (18 months−old mice compared to 8 months−old)
B
Age effects on protein abundance of CRLS members in liver
Stat sig
q−value > 0.1
Effect direction
Member protein up in older mice
Member protein down in older mice
−log10(p−value)
Age effect coefficients (18 months−old mice compared to 8 months−old)
Sex effects on protein abundance of CRLS members in liver
26
q−value < 0.1
Member protein up in male mice
RPL28
RPL7A
RPL21
RPL27
RPL34
RPLP2
Sex effect coefficients (females compared to males)
C
Age effects on protein abundance of CRLS members in lung
1
22
RPL3
RPL7
RPL15
RPL13
RPL19
RPL6
Sex effects on protein abundance of CRLS members in lung
Member protein up in female mice

**Figure S11. Complex-wide effects on the abundance of the cytoplasmic ribosomal large subunit (CRLS) differ between liver and lung.**

(A) Comparison of age-related differences in complex-wide correlation (y-axis) to complex-wide abundance (x-axis) of the CRLS across the 10 tissues. Liver and Lung are highlighted.

(B) Age- (left) and sex-related (right) abundance differences for proteins in liver, represented as volcano plots. The x-axis is the (left) age or (right) sex effect coefficient and the y-axis is the corresponding $-\log_{10}$(*p*-value). CRLS members highlighted with color based on direction of effect. Counts of proteins with differences are included (FDR < 0.1).

(C) Age- (left) and sex-related (right) abundance differences for proteins in lung, represented as volcano plots. The x-axis is the (left) age or (right) sex effect coefficient and the y-axis is the corresponding $-\log_{10}$(*p*-value). CRLS members highlighted with color based on direction of effect. Counts of proteins with differences are included (FDR < 0.1).
